# Modes of action and bio-fungicide potential of peptides derived from the bi-domain plant defensin MtDef5

**DOI:** 10.1101/2024.12.20.629707

**Authors:** Raviraj M. Kalunke, Ambika Pokhrel, Meenakshi Tetorya, Vishnu Sukumari Nath, Kirk J. Czymmek, Dilip M. Shah

## Abstract

*Medicago truncatula* bi-domain defensin MtDef5 exhibits antifungal activity at sub-micromolar concentrations against some fungal pathogens. It comprises two single-domain defensins, MtDef5A and MtDef5B, connected with a linker APKKVEP. MtDef5B is a more potent antifungal defensin than MtDef5A. We identified amino acid residues important for antifungal activity of MtDef5B and elucidated its modes of action (MoA). MtDef5B inhibited spore germination of *Botrytis cinerea (Bc)* at low micromolar concentrations. However, it did not inhibit spore germination of *Colletotrichum gloeosporioides (Cg)*. MtDef5B permeabilized the plasma membrane, induced reactive oxygen species and traveled to the nucleoli in germlings of *Bc*. Furthermore, a carboxy-terminal MtDef5A-derived GMA5AC peptide was selected for mutagenesis because of its lower cationicity than the corresponding MtDef5B-derived peptide. GMA5AC inhibited spore germination of *Bc*, but not of *Cg*. However, GMA5AC_V2, a variant of GMA5AC, inhibited spore germination of *Cg* and exhibited multi-faceted MoA. Spray-application of GMA5AC_V2 on the leaves of pepper plants demonstrated preventive and curative control of the gray mold disease. Furthermore, when applied topically on tomato fruits pre-inoculated with the pathogen *Cg*, this peptide reduced anthracnose disease symptoms. This study highlights the potential of short chain defensin-derived peptides for management of fungal diseases.

**Highlights:** The bi-domain MtDef5-derived antifungal peptides exhibit multiple modes of action and confer resistance against the gray mold and anthracnose diseases in pepper plants and tomato fruits, respectively.

## Introduction

Chemical fungicides are a vital arsenal in the crop protection toolkit (Steinberg and Gurr, 2020). Since they have a single biochemical target in fugal pathogens, their overuse has significantly accelerated resistance development (Hawkins and Fraaije, 2023). The ever-increasing spread of fungicide resistance has created an urgent need for development of functionally diverse fungicides with multi-faceted modes of action (MoA) that can minimize the risk of fungal resistance. Plant antimicrobial peptides (AMPs) with vast structural and sequence diversity hold potential as a novel class of bio-fungicides with efficacy and safety (Lobo and Boto, 2022; Ma *et al*., 2023; Rosa *et al*., 2022; Tang *et al*., 2023; van der Weerden *et al*., 2013).

The innate immune system of plants uses a broad range of AMPs as a first line of defense against invading pathogens. Defensins are the best studied plant AMPs having a small size of 45-54 amino acids and broad-spectrum antifungal activity (Cools *et al*., 2017; Parisi *et al*., 2019). Despite their sequence diversity, plant defensins have a highly conserved three-dimensional (3D) structure comprising one *α*-helix and three antiparallel *β*-strands that are connected by four disulfide bonds forming a cysteine-stabilized αβ (CSαβ) motif (Kovaleva *et al*., 2020). Because of their potent antifungal activity *in vitro* and *in planta* and stability at broad range of pH and temperature, defensins are considered as potential antifungal agents for use in agriculture (Cools *et al*., 2017; Kovaleva *et al*., 2020; Parisi *et al*., 2019). The major determinant of the antifungal activity of defensins resides in the carboxy-terminal γ-core motif (GXCX_3-9_C). Short length and low complexity of peptides containing the γ-core motif enable synthetic approaches to design more potent variants with varied MoA (Djami-Tchatchou *et al*., 2023; Tetorya *et al*., 2023). It is important to identify several short-chain defensin-derived peptides with high potency and different MoA. Different MoA of peptides will ensure effective management of fungal pathogens, minimize the risk of resistance development and offer opportunities for synergistic applications.

*Medicago truncatula* defensin 5 (MtDef5) is a bi-domain defensin containing two monomers, 5A and 5B, linked by a 7-amino acid peptide APKKVEP. Each monomer is 50-amino acids long differing from each other in eight residues. This bi-domain defensin carries a net charge of +16 and 39% hydrophobic amino acids (Islam *et al*., 2017). It exhibits broad-spectrum antifungal activity against *Bc*, *Fusarium* spp., *Alternaria brassicicola* and *Colletotrichum higginsianum* at sub-micromolar concentrations and exhibits multi-faceted MoA (Islam *et al*., 2017).

Herein, we strategically evaluated elements of the MtDef5 bi-domain peptide to optimize our understanding and potentially improve performance of the full-length and truncated variants as spray-on peptides. We determined *in vitro* antifungal activity of the single-domain MtDef5B defensin and its variants against multiple fungal pathogens and identified amino acids important for antifungal activity. MtDef5B was active against all pathogens tested except *Colletotrichum gloeosporioides (Cg)*. This defensin displayed multi-faceted MoA against *Bc*. The MtDef5A-derived short chain γ-core peptide GMA5AC was selected for generating variants because of its lower cationicity than the corresponding MtDef5B-derived peptide, and their antifungal activity against *Bc* and *Cg* was determined. One variant, GMA5AC_V2, displayed higher antifungal activity against these pathogens than GMA5AC. When applied topically on pepper plants and tomato fruits, GMA5AC_V2 was highly effective in reducing the gray mold and anthracnose disease symptoms, respectively.

## Materials and methods

### Growth of fungal cultures and preparation of spore suspension

The fungal strains of *Bc* T4, *Fusarium virguliforme (Fv)* NRRL 22292 *(Mont-1), Fg* PH-1, *Cg* 550 and an oomycete *Pc* were each grown in their respective growth media and conditions as indicated in Supplementary Table S1. *Bc* T-4 was a gift from Dr. M. Dickman at Texas A&M University. *Fg* and *Fv* isolates were obtained from the labs of Dr. J.-R. Xu at Purdue University and A. Fakhoury at Southern Illinois University, respectively. *Cg* was kindly provided by Dr. N. Peres, Gulf Coast Research and Education Center, University of Florida. *Pc* isolate was obtained from Dr. R. Grummet at Michigan State University. The zoospores of *Pc* were harvested as described previously (Mansfeld *et al*., 2020). Fungal spores were harvested by flooding the fungal growth plates with ∼5 ml of sterile water. Then, the spore suspension was filtered through two layers of Miracloth, centrifuged at 13,000 rpm for 1 min, washed three times using sterile water, and re-suspended in low-salt Synthetic Fungal Medium (SFM) (Liang *et al*., 2001). The spore suspension of each fungal pathogen was adjusted to the desired spore density using a hemocytometer.

### Recombinant expression of MtDef5B and its variants in *Pichia pastoris* and their purification

MtDef5B variants used in this study are shown in Table 1. Genes encoding these variants were generated using the QuickChange II site-directed mutagenesis kit (AGLS200523, Agilent Technologies, Santa Clara, CA). The synthetic codon-optimized genes encoding MtDef5B and its variants were obtained from GenScript (Piscataway, NJ). Ala codon was added to the 5’ end of each gene to obtain correct processing of each peptide in *P. pastoris*. Each synthetic gene was cloned into pPICZα-A integration vector in-frame with the *α*-mating factor secretion signal sequence containing KEX2 without the Glu-Ala repeats. The resulting recombinant plasmid, pPICZα-A containing each gene was linearized using the restriction enzyme *Sac*I or *Pme*I and then introduced into competent cells of the *P. pastoris* strain, X33, by MicroPulser (Bio-Rad, Hercules, CA). *P. pastoris* transformants for each gene were grown in the liquid medium for expression of each peptide as described (Li *et al*., 2019; Velivelli *et al*., 2020). Each peptide secreted into the medium was purified to homogeneity using Fast Protein Liquid Chromatography (FPLC, model ÄKTA Pure, Cytiva Marlborough, MA) and Reverse Phase-High Performance Liquid Chromatography (HPLC, Agilent Technologies, Santa Clara, CA) following the protocols described in (Islam *et al*., 2017; Li *et al*., 2019; Velivelli *et al*., 2020). HPLC purified peptides were lyophilized and re-suspended in the nuclease-free water. The peptide content was measured using NanoDrop 2000c spectrophotometry and Pierce BCA Protein Assay (Thermo Fisher Scientific, Grand Island, NY).

**Table 1:**
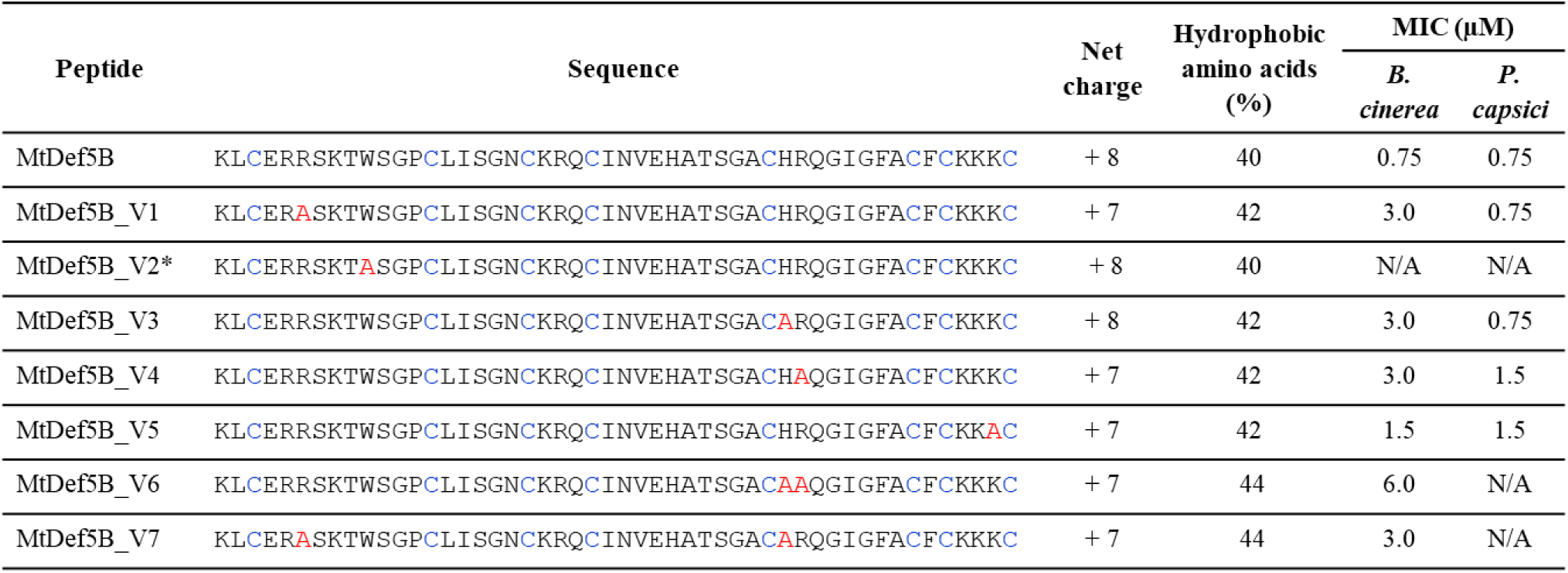
The deduced amino acid sequence, net charge, percentage (%) of hydrophobic amino acids and antifungal activity against plant fungal pathogens of MtDef5B and its variants. MtDef5B and its variants showed antifungal activity against *Bc* and an oomycete *Pc*. The resazurin-based fungal cell viability assay was used to determine the MIC values of peptides against *Fo* and *Pc.* Cysteine residues are highlighted in blue; Replaced amino acids in the original sequence of MtDef5B or GMA5AC are highlighted in red; MIC, minimum inhibitory concentration; N/A, not assessed. *MtDef5B_V2 could not be expressed in *P. pastoris*.

### Peptide synthesis and purification

The chemically synthesized peptides listed in Figure 1A were obtained from WatsonBio (Houston, TX). Each peptide with an initial purity of >90% was used without any further purification. The concentration of each peptide was determined using NanoDrop 2000c spectrophotometry and confirmed using Pierce BCA Protein Assay (Thermo Fisher Scientific, Grand Island, NY).

**Figure 1.**
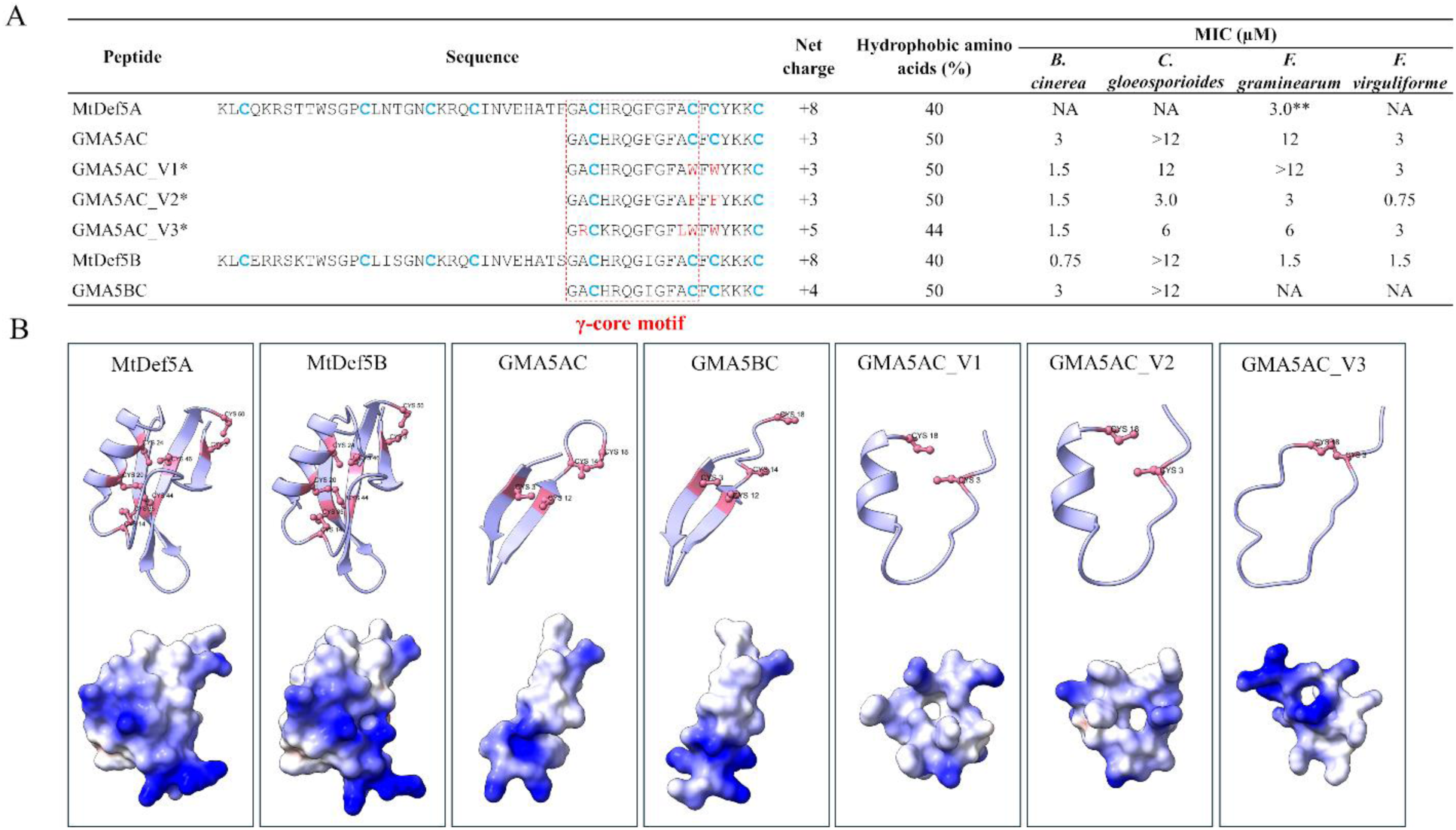
The amino acid sequences, properties, antifungal activity against plant fungal pathogens and structural predictions of MtDef5-derived peptides. (A) Amino acid sequences of MtDef5A, MtDef5B, GMA5AC, GMA5BC, GMA5AC_V1, GMA5AC_V2, and GMA5AC_V3 along with their net charge, percentage of hydrophobic amino acids and MIC against fungal pathogens. The γ-core motif is highlighted within the red box and the cysteine residues are shown in blue. The amino acid substitutions in the original sequence of MtDef5B or GMA5AC are highlighted in red; *C-terminal amidation modification; **MIC against *Fg* was reported previously by Islam et al. (2017); N/A, not assessed. (B) Structures of MtDef5A, MtDef5B, GMA5AC, GMA5BC, GMA5AC_V1, GMA5AC_V2, and GMA5AC_V3 as predicted by Alphafold2 (see Supplementary Table S2 for structural confidence scores). The NMR structure of MtDef4 (PDB: 2LR3) was used as a template. The predicted structures were visualized using ChimeraX. The upper panel shows the ribbon diagram where cysteine residues are highlighted in pink color. The potential disulfide atomic bonds are also highlighted in pink color with ball stick. The lower panel shows the electrostatic surface structures where blue color indicates the positively charged region and white surface indicates the region with zero net charge.

### Structure prediction and bioinformatics analysis

The structures of MtDef5A, MtDef5B, and their C-terminal variants were predicted using AlphaFold2 (Jumper *et al*., 2021) utilizing previously published NMR structure of MtDef4 (PDB: 2LR3) as a template (Sagaram *et al*., 2013). The top predicted structures were analyzed and visualized using UCSF Chimera X (Pettersen *et al*., 2021). The homologs of MtDef5B were searched using BLASTP against the non-redundant (nr) protein sequences database in NCBI (Camacho *et al*., 2009). The resulting output sequences were filtered manually based on their unique amino acid differences in the γ-core motif. Multiple sequence alignment of filtered homologs (Supplementary Fig. S1) was performed using Clustal Omega (Sievers and Higgins, 2018).

### Antifungal activity assays

*In vitro* antifungal assays to determine minimal inhibitory activity (MIC) of each peptide against fungal or oomycete pathogens were performed using the resazurin cell viability assay as described previously (Tetorya et al. 2023). Forty-five microliters of each peptide dilution (0.375, 0.75, 1.5, 3, 6, 12 μM) was added to each well of the microtiter plate containing 45 μL of (∼10^5^ spores/ml) spore suspension of fungal pathogens in SFM (Liang et al. 2001). The pathogen/peptide mixture was allowed to incubate for 48 h, then 0.01% (wt/vol) resazurin solution was added to each well and further incubated overnight. The change of the resazurin dye from blue color to pink or colorless indicated the presence of viable fungal cells. The MIC was determined as the lowest concentration of the peptide where no color change was seen. Microscopic examination was conducted to confirm these MIC values. Three biological and three technical replicates were conducted for determining *in vitro* antifungal activity of each peptide.

### Semi-*in planta* antifungal activity assays

The semi-*in planta* antifungal activity of MtDef5B and short chain peptides against *Bc* was determined using detached leaves from *N. benthamiana* plants as described previously (Li *et al*., 2019) with minor modifications. Briefly, detached pepper leaves were placed in Ziploc plastic containers maintaining high humidity. Then, 10 µL of spore suspension (∼5×10^4^ spores/mL) was placed carefully onto leaf samples, followed by 10 µL peptide or water inoculation at the same spot. Each peptide was tested at four different concentrations 0.75, 1.5, 3, and 6 µM against *Bc*. The disease lesions on the leaf were measured and white light and crop reporter images were taken 3 days after inoculation (DAI).

Organic tomato fruits, purchased from a supermarket, were used for drop-inoculation assays to test the ability of MtDef5B and GMA5AC_V2 to reduce anthracnose fruit rot symptoms caused by *Cg*. A minor wound on tomato fruit was generated using a sterile syringe, followed by the application of 10 μl of *Cg* conidia suspended in water (∼1×10⁵ spores/ml). After 14 hr, 10 μl of 24 μM of MtDef5B and GMA5AC_V2 (8x MIC) was applied. The inoculated tomato fruits were placed in Ziploc containers to maintain high humidity. Symptoms of anthracnose fruit rot were observed and photographed at 5-7 DAI.

### *In planta* antifungal activity assays

Four-week-old sweet pepper plants (*Capsicum annuum* cv. California Wonder) grown in the greenhouse were used for *in planta* spray application assays. Both preventive and curative *in planta* antifungal activity assays were performed. To test preventive *in planta* antifungal activity of peptides, pepper plants were first sprayed with 2 mL of water or peptide (6 µM). The sprayed plants were then air dried for 2 hr which was followed by inoculation with 2 mL of *Bc* spores (∼5×10^4^ spores/mL) suspended in 0.5x SFM media. To test curative *in planta* antifungal activity of peptides, pepper plants were first inoculated with *Bc* spores which was followed by water or peptide spray after 2 hr. The inoculated plants were transferred to Ziploc plastic containers under high humidity and disease symptoms were assessed after 3 days. The plants were scored based on six disease index categories: 0 – no symptoms, 1 = 1-25% disease, 2 = 26-50% disease, 3 = 51-75% disease, 4 = 76-95% disease, 5 = >96% disease or almost dead plant. Both white light as well as CropReporter (Phenovation B.V., Wageningen) images were taken for two representative leaves of each plant. The preventive and curative *in planta* antifungal activity assays were repeated at least twice with three technical replicates in each treatment.

### Internalization and subcellular localization of MtDef5B in *Bc*

Time-lapse laser scanning confocal microscopy was performed to monitor internalization and subcellular localization of the fluorescent DyLight550-labeled MtDef5B into *Bc*. The DyLight550 amine-reactive dye was used to label MtDef5B following the instructions supplied by the manufacturer (62263, Thermo Scientific). The *Bc* conidia (50 μL of ∼10^5^/mL) were placed in 10 mm microwell of 35 mm glass bottom microwell dishes (MatTek Corporation, Ashland, MA) and allowed to form germlings (10-12h later). Subsequently, DyLight550-MtDef5B (50 µL, 1.5 µM) and DNA specific staining dye Hoechst 33342 (final concentration: 0.1 μg/mL, H1399, Thermo Fisher Scientific, Grand Island, NY)) was added and Leica SP8-X confocal microscope was used for time-lapse imaging (15 min duration at 30 second intervals). DyLight550-MtDef5B was excited at 550 nm and fluorescence detected at 580–620 nm whereas Hoechst 33342 was excited at 405 nm and fluorescence detected at 415-460 nm, using a line switch strategy to prevent crosstalk. Simultaneously, bright-field images were captured using a transmitted light detector.

Subcellular localization of MtDef5B was further investigated by examining co-localization of DyLight550-MtDef5B, nuclear-staining dye Hoechst 33342, and nucleolus-specific RNA staining dye, Nucleolus Bright Green (NBG, final concentration: 1 μmol/L, N511-10, Dojindo Molecular Technologies, Japan) using super-resolution microscopy. *Bc* germlings were prepared for microscopy as described above for conidia. Germlings were incubated with DyLight550-MtDef5B (50 µL, 1.5 µM) and Hoechst 33342 (final concentration: 0.1 μg/mL) for 30 min before proceeding with microscopy. The co-localization of DyLight550-MtDef5B and Hoechst 33342 was visualized using a ZEISS Elyra 7 super-resolution microscope (ZEISS Microscopy, Jena, Germany) in lattice structured illumination microscopy mode at excitation and emission wavelengths of 561 and 570-620 nm (DyLight550), 405 and 420-480 nm (Hoechst 33342), 488 and 495-550 nm (NBG).

Single-or time-lapse z-stacks were acquired with a C-Apochromat 63X water immersion objective lens (Numerical aperture = 1.2), with a 63 nm x-y pixel size and 371 nm z-step size using leap mode taking 15 phases per image and a 30 ms exposure/frame. Images were visualized using Zen Black 3.0 SR FP2 software (ZEISS) and post-processed with Structure Illumination Mode (SIM^2^) with low contrast settings. The channel alignment was performed using standard fluorescent beads to maintain imaging precision (TetraSpeck TM microspheres, 0.1 μm, blue/green/orange/dark red, T7279, Thermo Fisher Scientific, Grand Island, NY).

### *Bc* membrane permeabilization assay using SYTOX Green (SG)

The nucleic acid staining fluorescent dye SG (Thermo-Fisher Scientific) was utilized to assess membrane permeabilization in *Bc* germlings induced by MtDef5B or short chain peptides following the method described by (Li et al. 2019). After exposure to MtDef5B (0.75 μM) or short chain peptides (1.5 μM) at their MIC concentration for 15 min in presence of 0.5 µM SG, *Bc* germlings were observed using Leica SP8-X confocal microscope (63X water immersion objective lens, HC PL Apochromat CS2), with excitation at 488 nm and emission between 510–598 nm and a z-stacking about 30 sec intervals.

### Induction of intracellular reactive oxygen species (ROS) upon peptide treatment

ROS induction in *Bc* germlings was analyzed after 10 min and 15 min exposure to MtDef5B or the short chain peptides. This analysis involved assessing intracellular induction using the ROS indicator dye 2′,7′-dichlorodihydrofluorescein diacetate (H_2_DCFDA) at a final concentration of 10 μM (Invitrogen, Carlsbad, CA) followed by fluorescence confocal microscopy. Each peptide was used at its MIC for 10 min and 15 min and ROS induction was observed using confocal microscopy (Leica SP8-X) with an excitation wavelength of 485 nm and an emission range of 575–649 nm. A z-stack at each time point was acquired using a 63X water immersion objective lens (HC PL Apochromat CS2) at 30 second intervals.

### *In vitro* protein translation inhibition assay

The luciferase gene mRNA was produced using *in vitro* transcription (RiboMAX Large Scale RNA Production System (Promega). The *in vitro* translation of the luciferase gene mRNA (200 ng) was performed using wheat germ extract system in presence of MtDef5B (Promega). A known eukaryotic cell translation inhibitor cycloheximide was used as a positive control and sterile water served as a negative control for the *in vitro* translation assay. The whole reaction mixture comprising luciferase mRNA, wheat germ extract system, amino acid mixture, RNase inhibitor (New England Biolabs) and different concentrations of MtDef5B (0.75, 1.5, 3.0, 6.0, 12, 24, 48 µM) or water or cycloheximide was incubated at 25°C for 1 hr. The Luciferase Assay Substrate, ONE-Glo (Promega) was added. Finally, the luciferase enzyme activity was measured using a microplate reader (Tecan Life Sciences) to reflect translation efficiency in the presence of different concentrations of MtDef5B. The percentage of translation inhibition was determined by comparing the luciferase activity in sterile water (negative control) to the luciferase activity in presence of various concentrations of MtDef5B.

### Gel retardation assay

Gel retardation assay procedure was adapted from (Velivelli *et al*., 2018) with minor modifications. The total RNA was extracted and purified from *Bc* mycelium grown in Potato Dextrose Broth (Difco) using the NucleoSpin RNA Plus Mini kit (Takara Bio) according to the manufacturer’s instructions. Genomic DNA contamination was removed using DNase I treatment (Thermo Fisher Scientific) as per manufacturer’s instructions. The luciferase mRNA was produced using RiboMAX Large Scale RNA Production Systems (Promega) as per manufacturer instructions.

The gel shift experiments were performed by mixing 200 ng of total RNA or 500 ng of luciferase mRNA with different concentrations of MtDef5B in 20 μl of binding buffer (10 mM Tris-HCl, pH 8.0, 1 mM EDTA). The reaction mixtures were incubated for 1 hr at room temperature and then mixed with 4 μl of 6x gel loading dye (New England Biolabs). Following electrophoresis in 1.2% agarose gel containing SYBR Safe DNA gel stain (Thermo Scientific) with 1x TAE buffer at 120 V for 45 min, the total RNA or luciferase mRNA-MtDef5B interaction was visualized using a Bio-Rad ChemDoc XRS+ system.

## Statistical Analysis

All experiments performed in this study consisted of two or more biological replicates when appropriate with at least three or more technical replicates. The statistical comparisons for disease lesion size in semi-*in planta* experiments and disease severity values in *in planta* experiments was performed using Wilcoxon test in *R*. Pairwise comparisons were made between peptide treated and no peptide treated controls, and the comparisons with *p*-value ≤ 0.05 were considered as statistically significant. The statistical comparisons of protein translation inhibition between treatment conditions were done using Analysis of variance (ANOVA) followed by a Tukey’s post hoc test. The comparisons with *p*-value ≤ 0.05 were considered as statistically significant.

## Results

### Antifungal activity of MtDef5B and its variants against *Bc*

MtDef5 is a bi-domain defensin consisting of MtDef5A and MtDef5B single domain defensins. Each domain consists of 50-amino acids with a net charge of +8 and 40% hydrophobic amino acids (Fig. 1A). The AlphaFold2 generated structures of MtDef5A and MtDef5B showed the presence of a similar cysteine-stabilized α/β motif comprising one α-helix stabilized through tetradisulfide array to three anti-parallel β-strands. The tetradisulfide array was Cys3-Cys50, Cys14-Cys35, Cys20-Cys44, and Cys24-Cys46 (Fig. 1B) typical for plant defensins (Islam *et al*., 2017).

We set out to characterize the antifungal activity and MoA of the single-domain MtDef5B defensin because of its previously reported slightly higher antifungal activity (MIC 1.0-1.5 µM) against *Fg* than that of MtDef5A (MIC 1.5-3.0 µM) (Islam *et al*., 2017). MtDef5B was successfully expressed in *P. pastoris*, purified to homogeneity and tested for antifungal activity. It inhibited the growth of fungal pathogens *Bc*, *Fg, Fv,* and an oomycete pathogen *Pc* with MIC values of 0.75 µM, 0 1.5 µM, 1.5 µM and 0.75 µM, respectively (Fig. 1A, Table 1). MtDef5B failed to inhibit spore germination of *Cg* and thus was not active even at the highest concentration of 12 µM tested. However, it did induce hyperbranching and shortening of fungal hyphae when compared with hyphal growth in absence of this peptide. These data revealed that even a single domain MtDef5B has broad-spectrum antifungal and antioomycete activity at low micromolar or sub-micromolar concentrations but retains some selectivity.

### MtDef5B single and double amino acid substitution variants exhibit reduced antifungal activity against *Bc*

To identify amino acids important for antifungal activity of MtDef5B, a series of variants was generated by substituting specific hydrophobic or cationic amino acids with Ala. These variants contained single or double amino acid substitutions. Amino acids selected for substitutions were located in the carboxy-terminal loop containing the γ-core motif and the amino-terminal loop known to be important for antifungal activity. Single amino acid substitution variants MtDef5B_V1 (MtDef5B^R6A^), MtDef5B_V2 (MtDef5B^W10A^), MtDef5B_V3 (MtDef5B^H36A^), MtDef5B_V4 (MtDef5B^R37A^) and MtDef5B_V5 (MtDef5B^K49A^) were introduced into *P. pastoris* for expression (Table 1). Despite repeated attempts, MtDef5B_V2 with a W10A substitution could not be expressed indicating its potential toxicity toward *P. pastoris*, lack of stability or secretion and thus could not be tested for antifungal activity. MtDef5B_V1, MtDef5B_V3, MtDef5B_V4 and MtDef5B_V5 were successfully expressed, purified and tested for antifungal activity against *Bc*. MtDef5B_V3 and MtDef5B_V4 variants consistently had the MIC value of 3 µM, whereas MtDef5B_V1 and MtDef5B_V5 had MIC values of 3 µM and 1.5 µM, respectively, as compared with the MIC value of 0.75 µM for MtDef5B. This analysis revealed the importance of R6, H36 and R37 for antifungal activity. Against the oomycete *Pc*, MtDef5B_V1 and MtDef5B_V3 exhibited the same potency as MtDef5B. However, MtDef5B_V4 and MtDef5B_V5 had two-fold reduction in the inhibitory activity against this pathogen as compared with MtDef5B.

To further confirm the importance of R6, H36 and R37, we generated MtDef5B variants containing two amino acid substitutions. Thus, MtDef5B_V6 (MtDef5B^H36A/R37A^) and MtDef5B_V7 (MtDef5B^R6A/H36A^) were expressed in *P. pastoris*, purified and tested for antifungal activity against *Bc* (Table 1). This analysis revealed that MtDef5B_V6 was consistently six-to eight-fold less potent than MtDef5B confirming the importance of H36 and R37 for antifungal activity of this defensin. Antifungal activity of MtDef5B_V7 was similar to that of the corresponding single mutants indicating no further loss of antifungal activity from combining the R6A and H36A mutations.

### Variants of the short-chain GMA5AC spanning the γ-core motif of MtDef5A exhibit potent antifungal activity against multiple fungal pathogens

The eighteen-amino acid short chain peptide GMA5AC (G33-C50) of MtDef5A has a net charge of +3 and 50% hydrophobic residues whereas the corresponding GMA5BC peptide (G33-C50) of MtDef5B has a net positive charge of +4 and 50% hydrophobic residues (Islam *et al*., 2017). These two peptides differ in only two amino acids. The F8 and Y15 in GMA5AC have been replaced with I8 and K15 in GMA5BC (Fig. 1A). Both short peptides inhibited the growth of *Bc* with an MIC of 3 µM. GMA5AC has the same MIC against *Bc* as GMA5BC despite having lower net charge. GMA5AC has scope to increase its net charge, so we selected this peptide for generating variants and testing their antifungal activity against *Bc*. Based on our previous work with GMA4CG_V6, a variant of MtDef4-derived seventeen-amino acid peptide GMA4CG spanning the γ-core motif (Tetorya *et al*., 2023), we designed two synthetic variants, designated GMA5AC_V1 and GMA5AC_V2, in which the two internal cysteine residues (C12 and C14) were replaced with either W or F, respectively, while retaining its net charge and percentage of hydrophobic residues. A third variant (GMA5AC_V3) was also synthesized which carried five amino acid replacements (A2R, H4K, A11L, C12W, C14W) relative to the sequence of GMA5AC resulting in the increased net charge from +3 to +5 and a concomitant decrease of hydrophobic residues from 50% to 44% (Fig. 1A).

The AlphaFold2-predicted structures of GMA5AC and GMA5BC comprised of two anti-parallel strands forming a β-sheet. In contrast, AlphaFold2-predicted structures of GMA5AC_V1 carrying CFC to WFW substitution and GMA5AC_V2 carrying CFC to FFF substitution resulted in the formation of only a single α-helix near the carboxy-terminus of each peptide (Fig. 1B). Interestingly, the structure of GMA5A_V3 having a net charge of +5 was essentially a random coil with one predicted C3-C18 disulfide bond. The predicted structural confidence and template modeling scores for peptides are shown in Supplementary Table S2.

We hypothesized that GMA5AC_V1 and GMA5AC_V2 with aromatic hydrophobic residues (W or F) substituting for the two internal cysteine residues will have more potent antifungal activity than GMA5AC (Tetorya *et al*., 2023). Indeed, GMA5AC_V1 and GMA5AC_V2 had two-fold higher antifungal activity (MIC 1.5 µM) against *Bc* than GMA5AC (MIC 3 µM) (Fig. 1). However, increasing the net charge and decreasing the percentage of hydrophobic residues did not result in a further increase in antifungal activity. Next, we tested antifungal activity of GMA5AC and its three variants against three different fungal pathogens. All three variants had more potent antifungal activity than GMA5AC against *Cg, Fg* and *Fv*. Against these pathogens, GMA5AC_V2 was at least four-fold more potent than GMA5AC (Fig. 1A). It was notable that GMA5AC had an MIC value of >12 µM against *Cg*, but GMA5AC_V2 had a MIC value of 3 µM (Fig. 1A). At 12 µM, all three variants of GMA5AC completely inhibited the growth of this pathogen, but MtDef5B, GMA5AC and GMA5BC were only partially effective (Supplementary Fig. S2). Thus, substitutions of C12 and C14 with F were more effective than with W in increasing the antifungal activity of the fungal pathogens tested.

### MtDef5B, GMA5AC and its variants reduce gray mold disease symptoms semi-*in planta*

We compared the semi-*in planta* antifungal activity of MtDef5B, GMA5AC and its three variants, and GMA5BC using the detached leaves of *N. benthamiana* as previously described (Li et al., 2019). Drops of freshly prepared spore suspension of *Bc* were applied onto each leaf, followed by drop application of each peptide or water on top. All peptides were tested for their semi-*in planta* antifungal activity at concentrations of 0.75, 1.5, 3 and 6 μM and the disease lesions were measured after 2-3 days of inoculation. At 0.75 μM, none of the peptides were effective in controlling the gray mold disease. However, at 1.5 μM, only MtDef5B was able to significantly (p ≤ 0.05) reduce disease lesions (Fig. 2). At 3 μM, we found that both GMA5AC and GMA5BC were ineffective in preventing the development of disease lesions. At this concentration, all three variants of GMA5AC were more effective in reducing disease lesions than GMA5AC. At 6 μM, all peptides (MtDef5B, GMA5AC, GMA5AC_V1, GMA5AC_V2, and GMA5AC_V3) except GMA5BC significantly (*p* ≤ 0.05) reduced the gray mold disease lesions. (Fig. 2).

**Figure 2.**
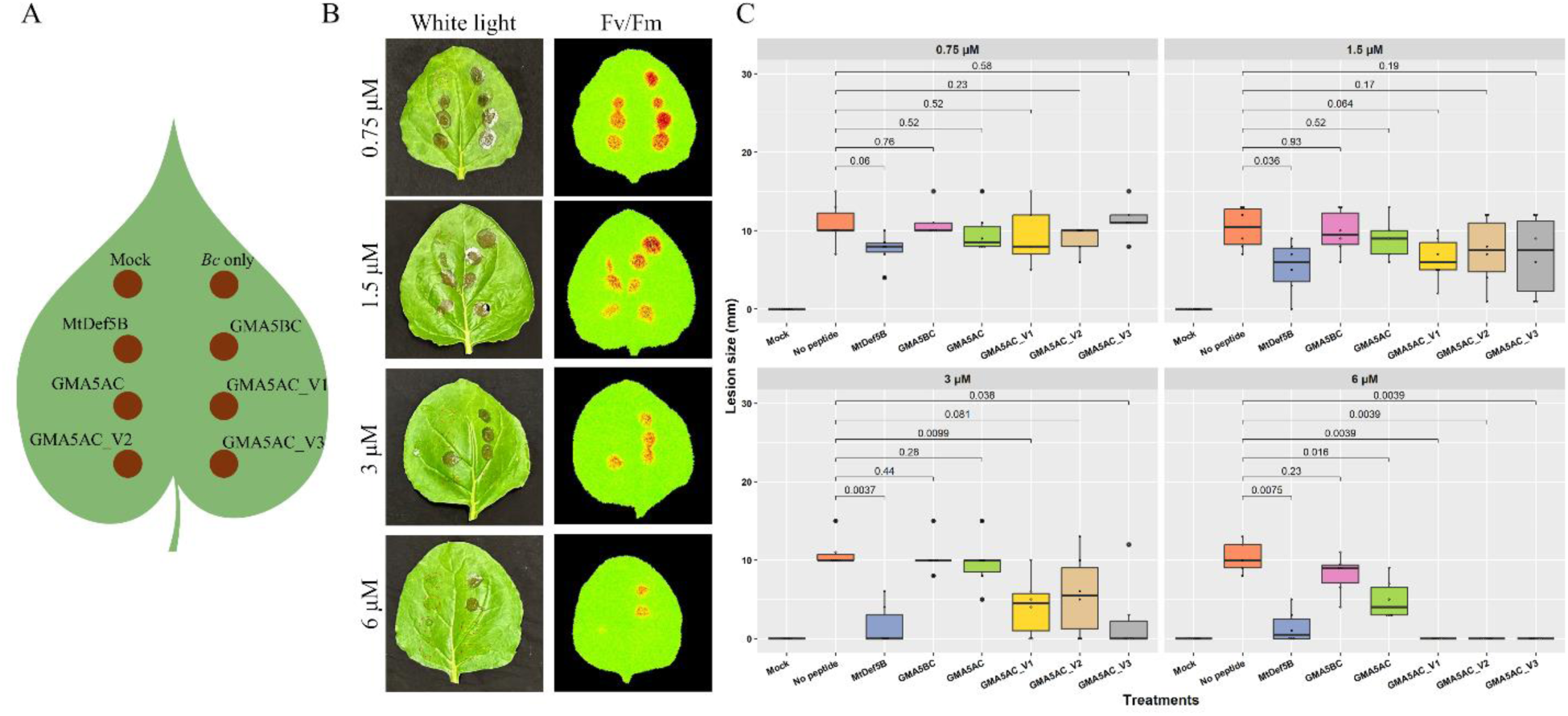
Topical application of MtDef5B, GMA5BC, GMA5AC, GMA5AC_V1, GMA5AC_V2, and GMA5AC_V3 on detached *N. benthamiana* leaves provides resistance against gray mold disease. (A) Scheme for application of MtDef5B, GMA5BC, GMA5AC, and GMA5AC_V1-V3 on detached leaves. **(B)** Gray mold disease lesions on *N. benthamiana* leaves were observed and photographed at 3 days after inoculation (DAI). The representative white light and CropRepoter (fv/fm) images of the leaves with each peptide tested at concentrations of 0.75 μM, 1.5 μM, 3 μM, and 6 μM are shown. Red color in CropReporter images represents extensive tissue damage and low photosynthetic efficiency (Fv/Fm) whereas green color indicates healthy tissue and higher photosynthetic efficiency. The spots with no disease are marked with red dashed circles for easier visualization (N=6 and R=2, where N refers to number of leaves and R refers to number of biological replicates). **(C)** Box plots showing the gray mold disease lesion size (in mm) in *N. benthamiana* leaves for each peptide treatment. The statistics were performed using Wilcoxon test and the *p*-values among the compared groups are indicated. *p*-value ≤ 0.05 is considered statistically significant.

### Spray-applied MtDef5B and GMA5AC_V2 are highly effective in providing preventive and curative control of gray mold disease in pepper plants

We tested MtDef5B, GMA5AC and its variants for their ability to confer preventive and curative antifungal activity against the gray mold disease on four-week-old pepper plants. For preventive antifungal activity, pepper plants were initially sprayed with MtDef5B, GMA5AC, GMA5AC_V1, GMA5AC_V2 and GMA5AC_V3 each at 6 μM followed by a spray-inoculation with *Bc* spores after a 2 hr interval. This concentration was chosen because under semi-*in planta* conditions all peptides were highly effective in preventing gray mold disease symptoms in detached *N. benthamiana* leaves. This preventive *in planta* antifungal assay revealed that pepper plants sprayed with MtDef5B, GMA5AC, and GMA5AC_V2 had significantly (*p* ≤ 0.05) reduced disease symptoms when compared to the no peptide control (Fig. 3). Interestingly, we found that GMA5AC_V2 provided much better protection against the disease when compared to GMA5AC, GMA5AC_V1 as well as GMA5AC_V3.

**Figure 3.**
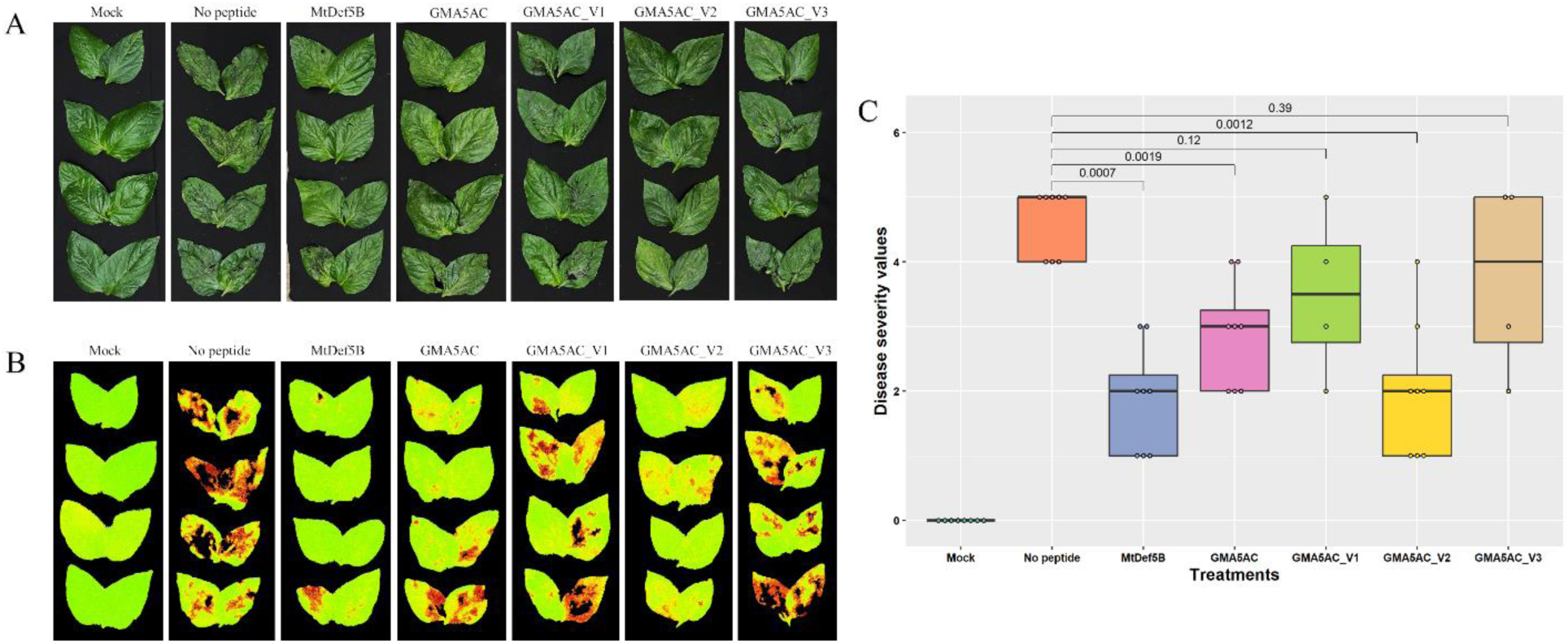
**MtDef5B, GMA5AC, and GMA5AC_V2 exhibit preventive antifungal activity against gray mold disease in pepper plants**. **(A)** White light and **(B)** CropReporter images of *in planta* preventive antifungal activity of MtDef5B and its variants used at a concentration of 6 μM against *B. cinerea*. The representative leaves of each plant and treatment were detached and imaged under normal light as well as on CropReporter. Red color in CropReporter images indicates extensive tissue damage and low photosynthetic efficiency (fv/fm) whereas green color indicates healthy tissue and higher photosynthetic efficiency (N=8 and R=2, where N refers to number of plants and R refers to number of biological replicates). **(C)** The box plot showing the disease severity values of gray mold disease on pepper plants. The disease rating was given based on a 0-5 scale, where 0 was a healthy plant with no symptoms and 5 indicated a severely diseased or dead plant. The statistics were performed using Wilcoxon test and the *p*-values among the compared groups are indicated. *p*-value ≤ 0.05 is considered statistically significant.

To evaluate curative antifungal activity of these peptides, pepper plants were first spray-inoculated with fungal spores. After two hr, pepper plants were spray-applied with each peptide at a concentration of 6 μM. Pepper plants sprayed with all peptides except GMA5AC_V3 provided significant (*p* ≤ 0.05) protection against gray mold disease when compared to the no peptide control (Fig. 4). Interestingly, GMA5AC_V2 outperformed all other peptides in providing curative protection against the gray mold disease under the conditions used (Fig. 4). Overall, our preventive and curative *in planta* antifungal assays revealed that GMA5AC_V2 offers greater protection against gray mold disease in pepper than other short peptides tested here.

**Figure 4.**
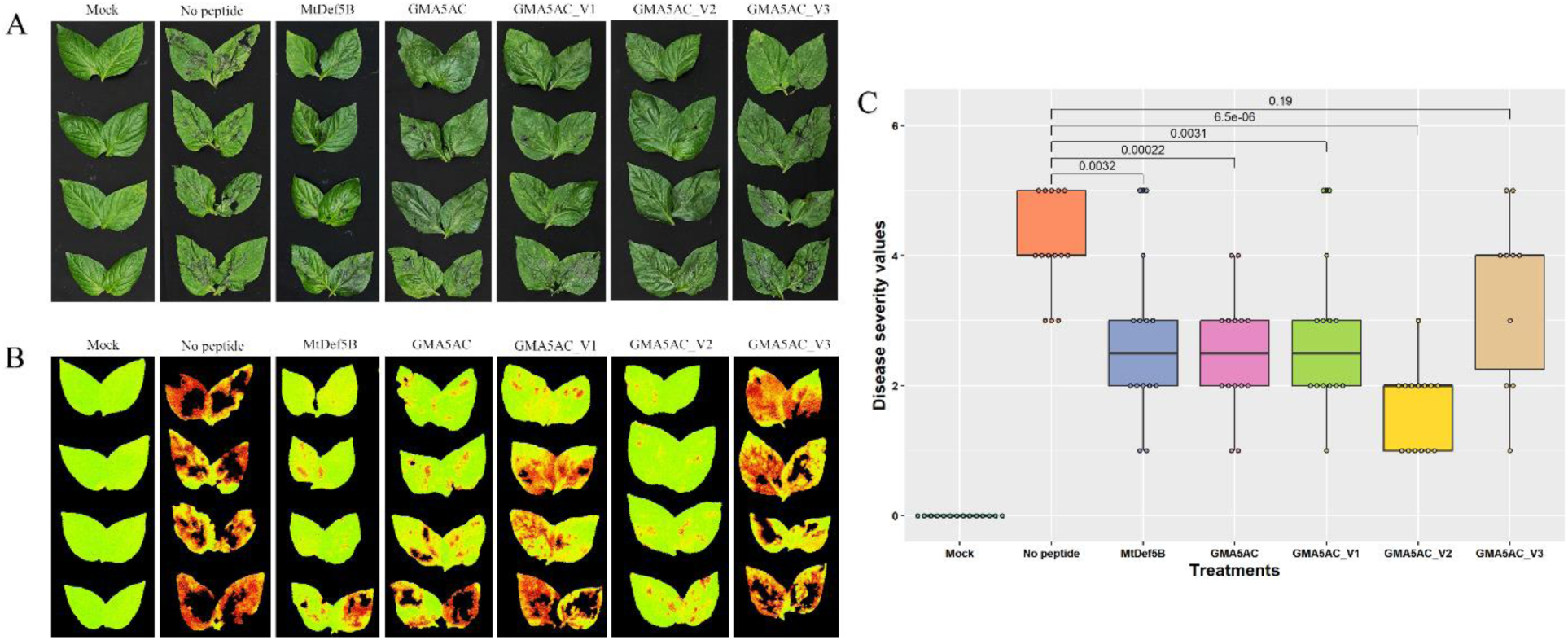
MtDef5B, GMA5AC, GMA5AC_V1, and GMA5AC_V2 exhibit curative antifungal activity against gray mold disease in pepper plants. **(A)** White light and **(B)** CropReporter images of *in planta* curative antifungal activity of MtDef5B, GMA5AC and its variants used at a concentration of 6 μM against *B. cinerea*. The representative leaves of each plant and treatment were detached and imaged with normal light as well as on CropReporter. Red color in CropReporter images represents extensive tissue damage and low photosynthetic efficiency (fv/fm) whereas green color represents healthy tissue and higher photosynthetic efficiency. (N=14 and R=3, where N refers to numbers of plants and R refers to number of biological replicates) **(C)** The box plot showing the disease severity values of gray mold disease on pepper plants. The disease rating was given based on a 0-5 scale, where 0 was a healthy plant with no symptoms and 5 indicated a severely diseased or dead plant. The statistics were performed using Wilcoxon test and the *p*-values among the compared groups are indicated. *p*-value ≤ 0.05 is considered statistically significant.

### Topical application of MtDef5B and GMA5AC_V2 on detached tomato fruits confers reduces anthracnose fruit rot symptoms

Among all peptides tested for *in vitro* antifungal activity against *Cg*, GMA5AC_V2 exhibited the most potent antifungal activity. We therefore tested the ability of this peptide to confer resistance to fruit rot caused by this pathogen. Tomato fruits were each wound-inoculated with 10 µl of 1x 10^5^ conidia of this pathogen. At 14 hr post-inoculation, GMA5AC_V2 and MtDef5 were each applied topically at a concentration of 24 µM. GMA5AC_V2 caused significant reduction in the anthracnose fruit rot symptoms as compared with MtDef5B indicating that a variant of the defensin-derived short peptide is more effective in controlling this disease than the full-length defensin (Fig. 5A, B).

**Figure 5.**
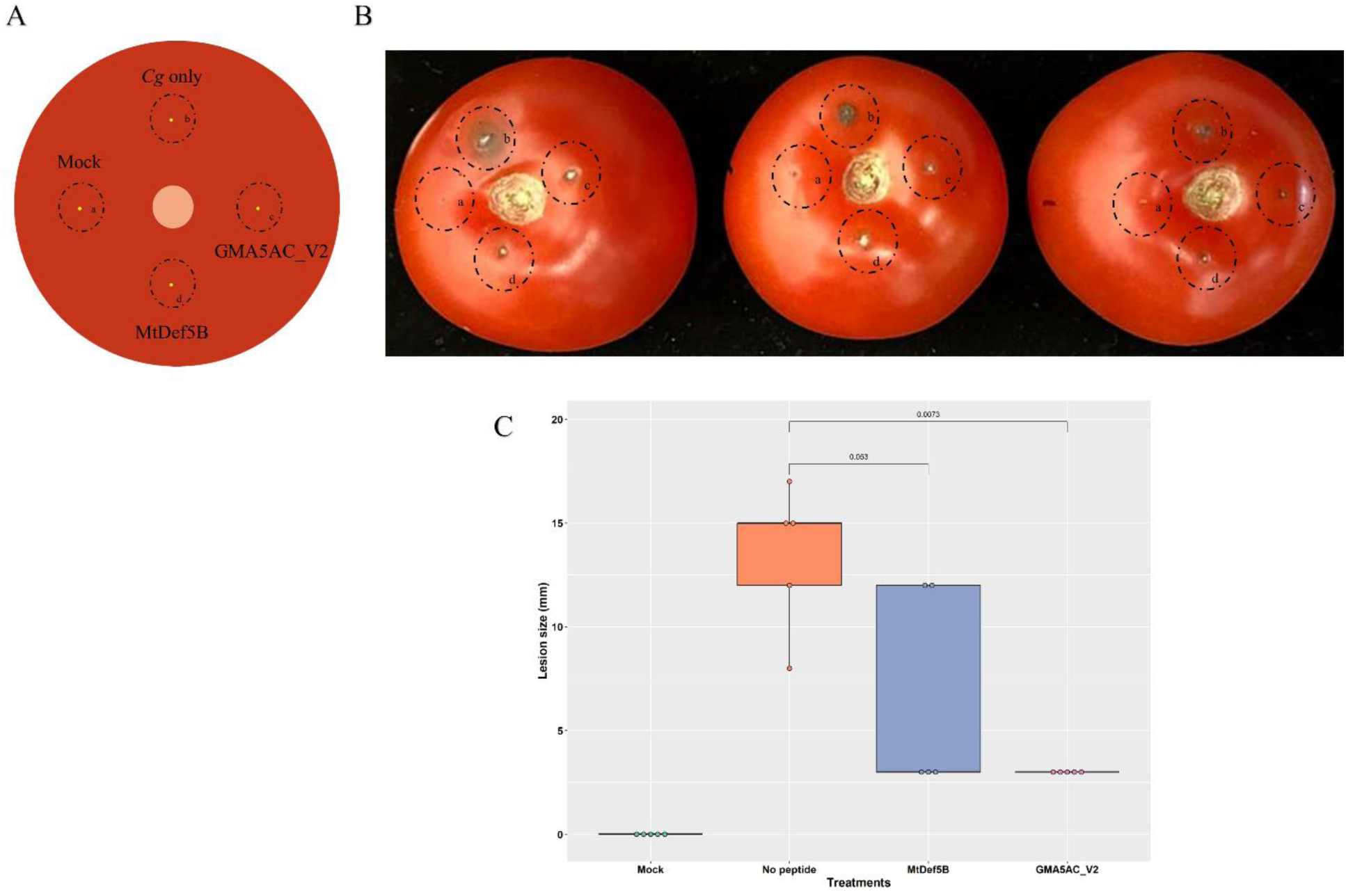
**Topical application of MtDef5B and GMA5AC_V2 on detached tomato fruits provides resistance against anthracnose fruit rot**. **(A)** Scheme for application of MtDef5B and GMA5AC_V2 peptide on the tomato fruit: A minor injury on each fruit was made using a sterile syringe, followed by the application of 10 μl of *C. gloeosporioides* conidial suspension (1 × 10⁵ conidia/ml). After 14 hr, 24 μM of each peptide was spray-applied (N=10 and R=2, where N refers to number of tomatoes and R refers to biological replicates). **(B)** Anthracnose fruit rot symptoms on tomato fruits were observed and photographed at 7 DAI, **(C)** The box plot showing the lesion size (mm) of anthracnose fruit rot on tomato fruits. The statistics were performed using Wilcoxon test and the *p*-values among the compared groups are indicated. *p*-value ≤ 0.05 is considered statistically significant.

### Intracellular uptake and subcellular localization of MtDef5B

As previously reported, the bi-domain MtDef5 was internalized by *Fg* and *Neurospora crassa (Nc)* co-localized with cellular membranes, then travelled to the nucleus and finally became dispersed in other subcellular locations (Islam *et al*., 2017). To determine if the uptake and subcellular localization of the single-domain MtDef5B were similar to that of the bi-domain MtDef5, MtDef5B was labeled with the fluorophore DyLight550 and its uptake was monitored by live-cell imaging of *Bc* germlings using confocal microscopy.

The DyLight550-MtDef5B has two-fold lower *in vitro* antifungal activity than the unlabeled MtDef5B against *Bc* (data not shown). Therefore, DyLight550-MtDef5B at MIC (1.5 µM) was added to *Bc* germlings and subjected to time-lapse imaging using confocal microscopy (Fig. 6A and 6B). Conspicuous cytoplasmic vacuolization was observed after two min of DyLight550-MtDef5B treatment and subsequently increasing in germlings. The DyLight550-MtDef5B initially bound to the surface of the germ tube tip while also creating entry points along the germ tube. A gradual MtDef5B uptake into germlings was observed over time. Within 15 min, the entire fungal hyphal structure became saturated with the DyLight550-labeled peptide. However, elevated accumulations of MtDef5B were observed in specific areas of the fungal cell (Fig. 6A). Next, we used the fluorescent nuclear-staining dye Hoechst 33342 to determine if the peptide travelled to the nuclei. The peptide entered germling cells at random entry sites (Fig. 6B, b, T= 6.42). At 15 min, co-localization of DyLight550-MtDef5B with Hoechst 33342 stained nuclei was observed (Fig. 6B). Further, we performed Structured Illumination Microscopy (SR-SIM) on *Bc* germlings treated with DyLight550-MtDef5B for 30 min (Fig. 6C). Again, MtDef5B peptide localized within Hoechst 33342 labeled nuclei of germling cells. Next, we applied the nucleolus-specific RNA staining dye, Nucleolus Bright Green (NBG), to determine if the peptide was localized in the nucleoli of the germling cells. We observed a very clear co-localization of the labeled peptide with NBG-stained nucleoli (Fig. 6C) indicating that the peptide targets nucleoli where ribosomal biogenesis takes place.

**Figure 6.**
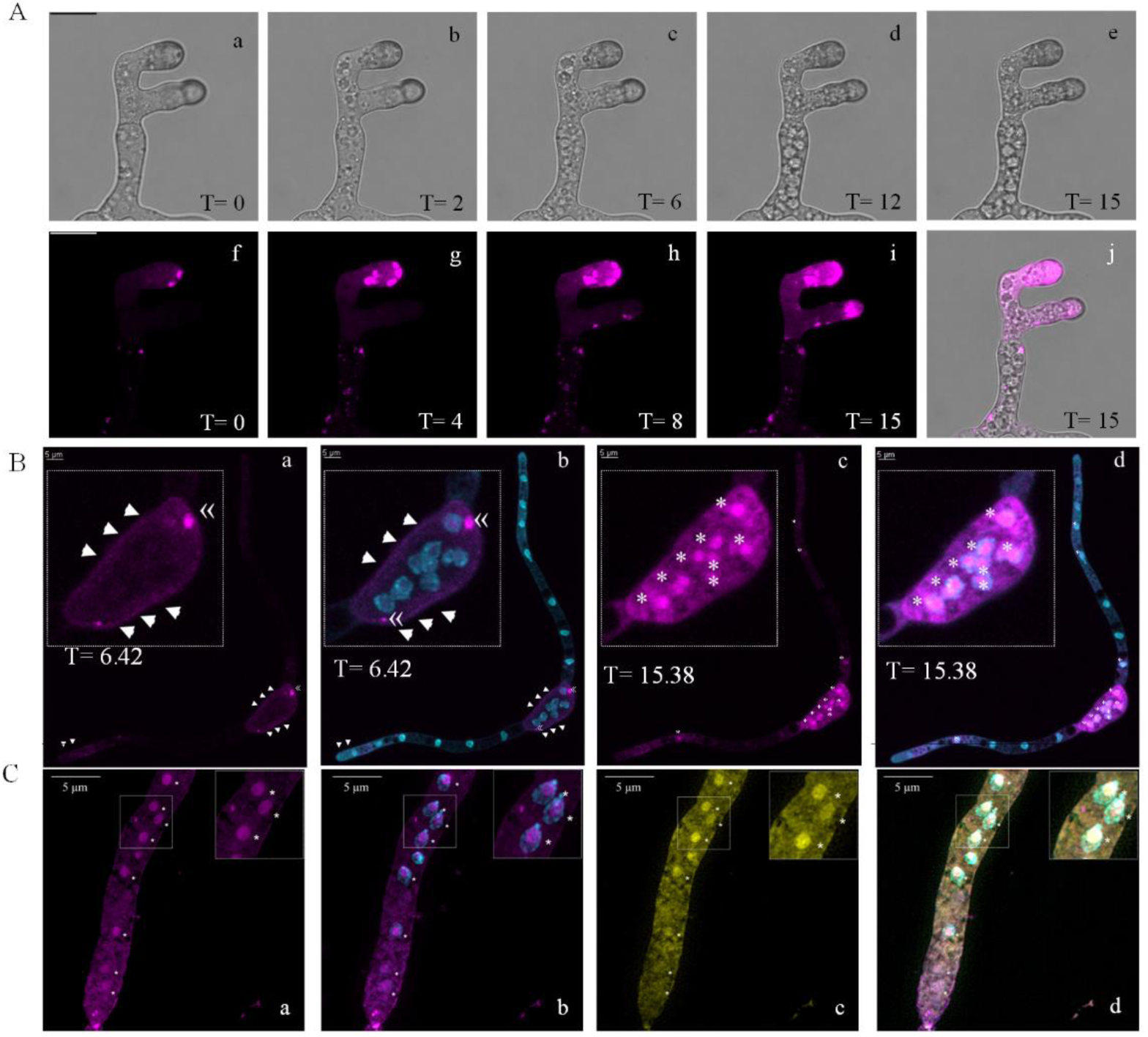
Confocal and super-resolution lattice structured illumination microscopy (SR-SIM) reveal internalization and subcellular localization of DyLight550-labeled MtDef5B in *B. cinerea* germlings. **A)** Time lapse confocal microscopy of *B. cinerea* germlings showing the internalization of DyLight550 labeled MtDef5B (magenta). Brightfield images of *B. cinerea* at T=0 (a), after 2 min (b, T=2). Early cytoplasmic vacuolization was observed. Subsequently increasing vacuolization in germlings was observed (c to e). DyLight550-MtDef5B initially bound to the surface of the hyphal tip while also creating entry points (f, T=0) and a gradual MtDef5B uptake was observed over time (g, h, i). Within 15 minutes, the entire fungal structure became saturated with DyLight550-labeled MtDef5B (i, j, T=15). However, a higher accumulation of MtDef5B was observed in specific areas of the fungal cell. Scale bar = 10μm. **B)** Time-lapse confocal microscopy of *B. cinerea* germlings showing internalization of DyLight550-MtDef5B and localization of fluorescence corresponding to nuclei stained with Hoechst 33342 (cyan). Binding of DyLight550-MtDef5B to fungal cell surface marked by arrows (a, T= 6.42) and peptide entry sites marked by guillemets (a, T= 6.42). These entry sites show separation from nuclear-staining dye Hoechst 33342 (b, T= 6.42). At 15 min, a higher concentration of DyLight550- MtDef5B accumulated within Hoechst 33342 labeled nuclei indicated by asterisk (c and d, T=15.38). **C)** SR-SIM images of *B. cinerea* germlings showing DyLight550-MtDef5B internalization after 30 min. A higher concentration of DyLight550-MtDef5B (a) was localized inside the Hoechst 33342 labeled nuclei and (b) indicated by asterisk. DyLight550-MtDef5B specifically co-localized with Nuclear Bright Green in nucleoli, the site of ribosomal biogenesis (c and d).

### MtDef5B disrupts the plasma membrane and rapidly induces reactive oxygen species (ROS) in *Bc*

We investigated whether single-domain MtDef5B was capable of permeabilizing the fungal plasma membrane. *Bc* germlings were treated with 0.75 µM MtDef5B for 15 min in presence of the dye SG (0.5 µM) and the uptake of the SG was observed using confocal microscopy. SG is a membrane-impermeant dye that fluoresces upon binding to RNA and DNA, serving as an effective indicator of membrane permeability. SG uptake was observed in nearly all *Bc* germlings treated with this peptide (15 min), but not in the control germlings (Supplementary Fig S3).

The induction of ROS was visualized using confocal microscopy with the ROS-specific indicator dye H_2_DCFDA (10 µM). H_2_DCFDA is a non-fluorescent derivative of fluorescein (its reduced acetylated form) that becomes fluorescent only after the cleavage of its acetyl groups and subsequent oxidation within the cell, converting it to 2’,7’-dichlorofluorescein (DCF). ROS accumulation was observed in the hyphae of *Bc* germlings within 10 min of exposure to 0.75 µM MtDef5B (Supplementary Fig S4). The accumulation of ROS continued to be evident even after 15 min. The observed fluorescence indicates that ROS-mediated mechanisms may contribute to the apoptotic or necrotic death of fungal cells induced by MtDef5B (Supplementary Fig S4).

### MtDef5B inhibits protein translation *in vitro*

Since MtDef5B is targeted to the nucleolus, we surmised that it might interfere with protein translation as part of its MoA. We tested the potential of MtDef5B to inhibit protein translation using a eukaryotic wheat germ extract *in vitro* translation system for the luciferase gene. *In vitro* translation of the mRNA was performed using wheat germ extract system in presence of different concentrations of MtDef5B ranging from 0.75 to 48 µM. Cycloheximide, a eukaryotic protein translation inhibitor, was used as a positive control and sterile water served as a negative control for the *in vitro* translation assay. The luciferase enzyme activity was measured using the microplate reader to reflect translation efficiency in the presence of MtDef5B. As expected, high concentration of the luciferase activity was detected in presence of sterile water whereas no luciferase activity was detected in presence of cycloheximide (Fig. 7A). MtDef5B at 0.75 µM reduced translation efficiency ∼8.0%. Increasing the concentration of the peptide to 48 µM drastically reduced translation efficiency ∼90% as compared to the negative control (sterile water), suggesting that MtDef5B inhibits protein translation process in a concentration dependent manner (Fig. 7A). We asked if MtDef5B’s ability to inhibit translation was due to its interaction with the ribosomal RNA (rRNA) of *Bc* and/or to luciferase messenger RNA. An electrophoretic gel mobility shift assay was used to evaluate binding of MtDef5B *to Bc* ribosomal RNA (Fig. 7B). Like another plant defensin MtDef4 (Li *et al*., 2024), MtDef5B showed little or no binding with rRNA, but MtDef5B_V6 (MtDef5B^H36A, R37A^) and MtDef5B_V7 (MtDef5B^R6A, H36A^) showed partial binding with rRNA at a high concentration of 12 µM. We also tested binding of MtDef5B and MtDef4 at various concentrations to luciferase mRNA. MtDef5B and MtDef4 showed little or no binding to luciferase mRNA (Fig.7C). These results suggest that MtDef5B inhibits protein translation *in vitro* not by binding to rRNA but perhaps by binding to ribosomal or other proteins playing a role in translation initiation, elongation or termination.

**Figure 7.**
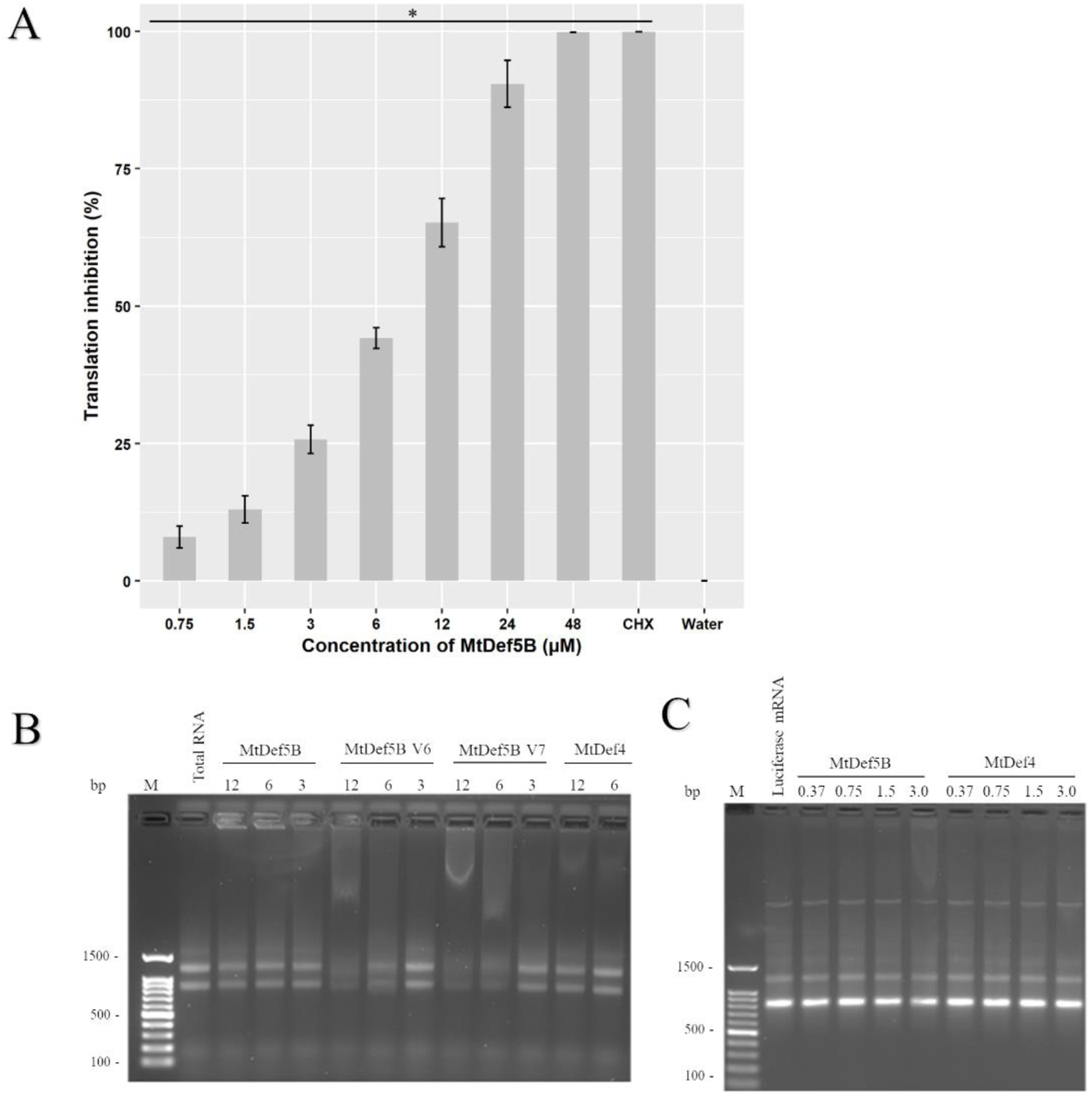
MtDef5B inhibits *in vitro* protein translation but shows minimal interaction with *B. cinerea* ribosomal RNA (rRNA) and luciferase mRNA *in-vitro*. A) *In vitro* translation assay confirmed the ability of MtDef5B to inhibit protein translation. The luciferase mRNA was produced using the RiboMAX™ Large Scale RNA Production System (Promega) and then translated to luciferase enzyme using the wheat germ extract. At the end, the luciferase activity was quantified to reflect the translation efficiency in presence of various concentrations of MtDef5B (0 to 48 μM). Translation inhibition percentage (%) was measured by comparing the luciferase activity in sterile water (negative control) to the luciferase activity in presence of various concentrations of MtDef5B. Sterile water was used as a negative control and cycloheximide was used as a positive control for translation inhibition. *Indicates significant differences between data of the negative control and the treated samples with *p* ≤ 0.05, based on Analysis of variance (ANOVA) and followed by a Tukey’s post hoc test. **B)** Electrophoretic gel mobility shift assay revealed minimal interaction of MtDef5B with *B. cinerea* total RNA. Different concentrations of MtDef5B (0 to 12 μM) were incubated with 200 ng total RNA extracted from *B. cinerea*. The total RNA incubated with sterile water was used as a control. **C)** Electrophoretic gel mobility shift assay revealed no interaction of MtDef5B with Luciferase mRNA. Different concentrations of MtDef5B or MtDef4 (0 to 3 μM) were incubated with Luciferase mRNA. The luciferase mRNA incubated with sterile water or MtDef4 (Li, et al., 2024) was used as a control. M, 100 bp DNA ladder.

### Short-chain variants of GMA5AC disrupt the plasma membrane, induce ROS production in *Bc* and inhibit protein translation *in vitro*

To gain insight into the MoA of the short-chain GMA5C_V2, we sought to determine the intracellular uptake and subcellular location of the peptide. However, the tetramethyl rhodamine (TMR)-labeled chemically synthesized peptide could not be dissolved in water and only partially in DMSO making it difficult to determine its antifungal activity and to perform subsequent confocal microscopy.

We conducted an SG uptake assay to assess the membrane permeabilizing ability of GMA4AC_V2. Within 15 min, nearly all *Bc* germlings treated with the peptide GMA5C_V2 at a concentration of 1.5 µM showed SG uptake indicating its ability to disrupt the fungal plasma membrane (Supplementary Fig. S5). *Bc* germlings challenged with GMA5C_V2 also accumulated ROS within 10-15 min (Supplementary Fig. S6). MtDef5B inhibited protein translation ∼80% inhibition at a concentration of 12 µM (Fig. 8). At this concentration, three variants of GMA5C inhibited *in vitro* translation ∼60%. However, both GMA5AC and GMA5BC truncated wild-type peptides showed lower translation inhibition when compared to the variant peptides (Fig. 8).

**Figure 8.**
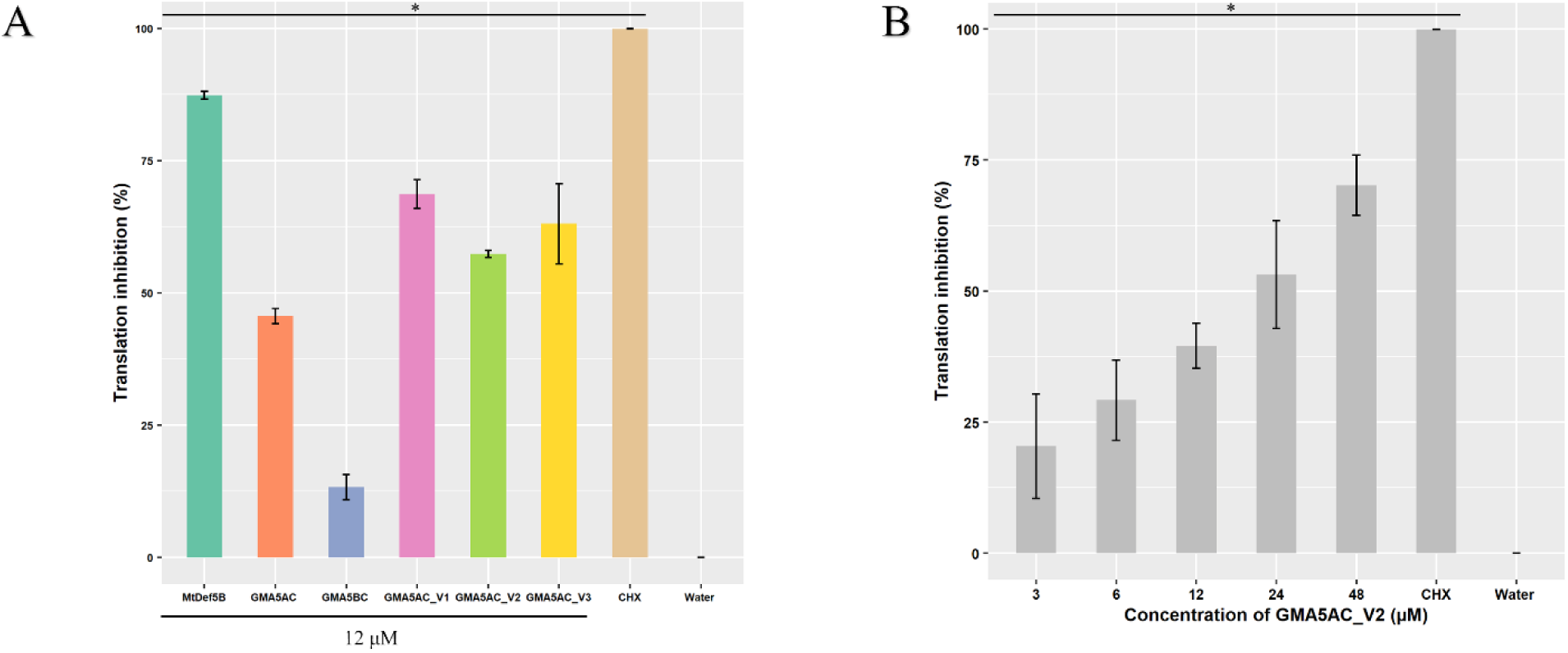
The *in vitro* translation assay confirmed the ability of GMA5AC_V1, GMA5AC_V2, and GMA5AC_V3 to inhibit protein translation. The luciferase gene was transcribed using the RiboMAX™ Large Scale RNA Production System (Promega) and then translated to luciferase enzyme using the wheat germ extract. At the end, the luciferase was quantified to reflect the translation efficiency in presence of **(A)** GMA5AC, GMA5BC, GMA5AC_V1, GMA5AC_V2, and GMA5AC_V3 (12 μM) and **(B)** in presence of GMA5AC_V2 (0 to 48 μM). The percentage (%) of translation inhibition was determined by comparing the luciferase activity in sterile water (negative control) to the luciferase activity in the presence of various concentrations of MtDef5B. Sterile water was used as a negative control and cycloheximide (CHX) was used as a positive control for translation inhibition. *Indicates significant differences between data of the negative control and the treated samples with *p* ≤ 0.05, based on Analysis of variance (ANOVA) and followed by a Tukey’s post hoc test.

Among all peptides tested, GMA5AC_V2 was the most effective in inhibiting the growth of *Cg* with an MIC of 3 µM. We tested the ability of GMA5AC_V2 to disrupt the plasma membrane and induce ROS production in this pathogen. SG uptake was observed in nearly all *Cg* germlings treated with this peptide (15 min, 3.0 µM), but not in the control germlings (Supplementary Fig. S7). ROS accumulation was observed in the hyphae of germlings within 10 min of exposure to 3 µM GMA5AC_V2 (Supplementary Fig. S8). The accumulation of ROS continued to be evident even after 15 min in peptide treated germlings, but not in the control germlings. Thus, the MoA of GMA5AC_V2 had some overlap with that of MtDef5B defensin involving plasma membrane disruption, ROS induction and *in vitro* protein translation inhibition, but details of its subcellular localization, inhibition of protein translation *in vivo* and interaction with other intracellular targets are not yet known.

## Discussion

In this study, we examined the potential as biofungicides of the bi-domain MtDef5 derived short chain peptides including MtDef5B domain and short chain γ-core peptides derived the carboxy-terminus of the MtDef5A domain. We characterized their antifungal activity against the economically important pathogens of crops and partially elucidated their MoA.

MtDef5B exhibited potent broad-spectrum inhibitory activity against multiple fungal pathogens including *Bc*, *Fg*, *Fv*, and an oomycete *Pc* with MICs ranging from 0.75 to 1.5 µM. However, it did not inhibit the growth of *Cg* at the highest concentration of 12 µM used in the assay indicating some selectivity in its antifungal action. *Cg* is an economically important pathogen that causes widespread postharvest infections in tropical fruits. Within the phylum Ascomycota, it belongs to a different family *Glomerellaceae* than *Bc*, *Fg* or *Fv*. It is likely that cell wall binding sites are either missing or have poor affinity for MtDef5B.

In the bi-domain MtDef5 defensin, we previously identified His36 and Arg37 residues in the A domain and His93 and Arg94 residues in the B domain as being essential for its antifungal activity. Through further site-directed mutagenesis of MtDef5B, we confirmed the importance of these two residues in the γ-core motif of this defensin for antifungal activity against *Bc*. In addition, Arg6 residue located in the N-terminal loop of this defensin also contributed to the antifungal activity. It is striking, however, that these three residues individually did not seem to be critical for the antifungal activity of this defensin against *Pc*. These findings suggest that other, as yet unidentified residues, are perhaps required for antifungal activity against these two pathogens in the amino acid sequence of MtDef5B. More extensive mutagenesis of the amino acid sequence is needed to fully elucidate specificity in the structure-activity relationships of this defensin against different pathogens.

We determined antifungal activity of the short chain GMA5AC peptide containing the carboxy-terminal 18 residues of the MtDef5A domain and found that it inhibited the growth of *Bc* and *Fv* with an MIC value of 3 µM. However, it was much less active against *Fg* having an MIC value of 12 µM. Like MtDef5B, it also failed to inhibit the growth of *Cg* at tested concentrations. We have previously reported efficacy of the variant of MtDef4 defensin-derived peptide spanning the active γ-core motif to confer curative as well as preventative resistance to *Bc* in *N. benthamiana* and tomato plants as a spray-on biofungicide (Tetorya *et al*., 2023). In this peptide, the conserved CFC motif was replaced by WFW motif resulting in enhanced *in vitro* as well as *in planta* antifungal activity. In this study, we have now extended these observations to variants GMA5AC_V1 and GMA5AC_V2 of the MtDef5A-derived peptide. When compared with GMA5AC, GMA5AC_V1 containing the WFW and GMA5AC_V2 containing the FFF motif in place of CFC motif exhibited two-fold more potent antifungal activity against *Bc*. Thus, the presence of aromatic strongly hydrophobic amino acids in specific locations of the short defensin-derived γ-core peptides seemed to be important for boosting their antifungal activity. It will be interesting to test if these observations hold true for defensin-derived γ-core peptides from other sequence-divergent defensins.

For decades, the control of fungal pathogens has relied heavily on chemical fungicides which inhibit very specific molecular targets. Although highly effective, this approach has led to multiple cases of fungicide resistance. Therefore, present-day research in crop protection is driven by the urgent need for sustainable fungicides that are effective, have a reduced risk of resistance, are eco-friendly, and non-phytotoxic. In semi-*in planta* studies using the detached leaves of *N. benthamiana*, we found that all peptides completely inhibited the development of gray mold disease lesions at 6 µM concentration. The concentration of peptides required to control *Bc* infection was 8X higher than the *in vitro* MIC which could be due to the tritrophic interaction happening between waxy cuticle of the leaf surface, peptide, and the fungus in the semi-*in planta* and *in planta* assays. The spray applied MtDef5B and GMA5AC_V2 provided excellent preventive as well as curative protection against the gray mold disease in pepper. The difference in preventive and curative protection provided by the spray-applied GMA5AC_V1 could be related to several factors. Even though the preventive protection provided by GMA5AC_V1 against *Bc* infection was not statistically significant, disease severity was relatively lower when compared to the no peptide control which clearly showed that this peptide is still effective in controlling the gray mold disease *in planta*. Further, among all the peptides tested for *in planta* antifungal activity in our study, we found that the truncated GMA5AC_V2 was effective in mitigating gray mold disease in pepper plants when compared to GMA5AC, suggesting that amino acid substitutions introduced in GMA5AC_V2 are important for its *in vitro* as well as *in planta* antifungal activity. While defensin derived peptides are advantageous due to their small size, simple structure, and modifiability, their transition to fungicidal commercial products depends on pioneering advancements in large-scale peptide manufacturing to make them financially viable.

A suite of potent antifungal peptides that exhibit different multi-faceted MoA has promise for new safe and sustainable innovative disease control tools. We have found that MtDef5B not only exhibited potent antifungal activity but also exhibited multi-element MoA in *Bc*. Like the parental bi-domain MtDef5 and other defensins we have studied (Islam *et al*., 2017; Li *et al*., 2024; Li *et al*., 2019), MtDef5B defensin rapidly disrupted the plasma membrane of this fungus, traveled to the nucleus and to other intracellular locations very rapidly. The fluorescently labeled MtDef5B localized to discrete spots in the fungal cell wall and entered the cell within 3 min of peptide challenge. There was a gradual increase of entry sites in hyphal tip area during the first 6 min of challenge and the peptide accumulated in the whole interior of the germling cell within min. Thus, entry of MtDef5B into fungal cells is more rapid than that of the bi-domain MtDef5 (Islam, et al. 2017).

Further subcellular localization studies reported here revealed that this defensin is also specifically localized into cytoplasm and nucleolus where it could potentially bind to ribosomes and/or ribosomal RNA. When incubated with total fungal RNA, MtDef5B revealed very little binding to rRNA as revealed by the gel retardation assay. This is similar to MtDef4 which showed binding to fungal ribosomes, but not to rRNA (Li *et al*., 2024). Both defensins, however, inhibit protein translation *in vitro*. The subcellular localization in the cytoplasm and nucleolus and/or inhibition of protein synthesis are emerging as conserved properties of plant antimicrobial peptides (Farkas *et al*., 2014; Li *et al*., 2024; Velivelli *et al*., 2020). It remains to be determined if protein translation inhibition by MtDef5B involves binding specific ribosomal subunits or to ribosomal proteins. Further, it will be of interest to determine if MtDefB inhibits protein translation in cells of *Bc* as part of its MoA.

Most cationic plant defensins contain in their sequences positively charged amphiphilic γ-core motif that acts as a major determinant of their antifungal activity. The carboxy-terminal peptide sequences containing the γ-core motif of these defensins exhibit antifungal activity (Slezina *et al*., 2022; Slezina *et al*., 2021). The sequence comparison of the amino acid sequences of 17 MtDef5B homologs from other plant species revealed that the γ-core motif sequence of each homolog is highly conserved but carries 1-3 amino acid substitutions relative to the sequence of the MtDef5B γ-core motif (Supplementary Fig S1). The γ-core peptide sequences of these defensins are potential candidates for development of as novel peptide-based fungicides for use in agriculture.

## Supplementary data

**Supplementary Fig. S1.** Amino acid sequence alignment of MtDef5B homologs from different plants. MtDef5B homologs sequences were obtained from NCBI and aligned using Clustal Omega. The *γ*-core motif is highlighted within the red box and the amino acid differences in the γ-core motif between the homologs are indicated in blue letters.

**Supplementary Fig. S2.** Representative microscopy images of *Colletotrichum gloeosporioides* treated with 12 μM MtDef5B, GMA5AC, GMA5BC, GMA5C_V1, GMA5C_V2, and GMA5C_V3. Images were taken 24 hr after incubation with peptides. Scale bar = 100 μm.

**Supplementary Fig. S3.** The MtDef5B permeabilizes the fungal plasma membrane in *B. cinerea* germlings. The fluorescence confocal microscopy images show SYTOX Green (SG) binding to the nuclei following treatment with 0.75 μM of MtDef5B in the presence of 0.5 μM SG after 15 min. Scale bar = 20 μm.

**Supplementary Fig. S4**. MtDef5B induces accumulation of Reactive Oxygen Species (ROS) in *B. cinerea* germlings. Confocal microscopy images showing ROS production (indicated by green fluorescence from H_2_DCFDA) in *B. cinerea* germlings after treatment with 0.75 μM MtDef5B for 10 and 15 min, Scale bar = 20 μm.

**Supplementary Fig. S5.** GMA5AC variants GMA5AC_V1, GMA5AC_V2, and GMA5AC_V3 permeabilize the fungal plasma membrane in *B. cinerea* germlings. The fluorescence confocal microscopy images show SYTOX Green (SG) binding to the nuclei following treatment with each peptide at 1.5 μM in the presence of 0.5 μM SG after 15 min. Scale bar = 20 μm.

**Supplementary Fig. S6.** GMA5AC variants GMA5AC_V1, GMA5AC_V2, and GMA5AC_V3 induce accumulation of Reactive Oxygen Species (ROS) in *B. cinerea* germlings. The confocal microscopy images showing ROS production (indicated by green fluorescence from H_2_DCFDA, 10 µM) in *B. cinerea* germlings after treatment with each peptide 1.5 μM for 10 and 15 min. Scale bar = 20 μm.

**Supplementary Fig. S7.** GMA5AC_V2 permeabilizes the fungal plasma membrane of *C. gloeosporioides* germlings. The fluorescence confocal microscopy images show SYTOX Green (SG) binding to the nuclei following treatment with 3.0 μm of GMA5AC_V2 in presence of 0.5 μM SG after 15 min. Scale bar = 20 μm.

**Supplementary Fig. S8.** GMA5AC_V2 induces accumulation of Reactive Oxygen Species (ROS) in *C. gloeosporioides* germlings. The confocal microscopy images showing ROS production (indicated by green fluorescence from H_2_DCFDA, 10 µM) in *C. gloeosporioides* germlings after treatment with 3.0 μM peptide for 10 and 15 min. Scale bar = 20 μm.

**Supplementary Table S1**: Growth media and conditions for culturing the fungi used in this study.

**Supplementary Table S2**: AlphaFold 2.0 predicted structural confidence and template modeling scores for selected peptides. The table compares various peptides based on the Predicted Local Distance Difference Test (pLDDT) and Predicted Template Modeling (pTM) scores.

## Acknowledgements and Funding

This work received partial support from the National Science Foundation-EAGER grant IOS:1955461, awarded to D.S. We also acknowledge the Advanced Bioimaging Laboratory (RRID: SCR_018951) at the Donald Danforth Plant Science Center (DDPSC) for imaging support, and the use of the Leica SP8-X confocal microscope, funded by an NSF Major Research Instrumentation grant (DBI-1337680) and the ZEISS Elyra 7 Super-Resolution Microscope funded by an NSF Major Research Instrumentation grant (DBI-2018962).

## Author contributions

Conceptualization, D.M.S; Methodology, R.M.K, A.P, M.T, and V.S.N; Formal analysis and investigation, R.M.K and A.P; Writing—original draft, R.M.K and A.P; Writing—review & editing, R.M.K, A.P, D.M.S, and C.J.K; Supervision, D.M.S and C.J.K.; Funding acquisition, D.M.S.

## Data Availability

The supporting data related to the findings of this study are included in the supplementary material of this article and linked raw data are available upon request.

