## Supplementary Materials for "Modes of action and bio-fungicide potential of peptides derived from the bi-domain plant defensin MtDef5"

| Plant species | Name/Accession# | Sequence Alignment |
|---|---|---|
| *Medicago truncatula* | MtDef5B | KLCERRSKTWSGPCLISGNCKRQCINVEH--ATSGACHRQGIGFACFCKKKC |
| *Solanum tuberosum* | XP_015160567.1 | RVCERRSSTWSGPCFDTGNCNRQCINWEH--ASSGACHREGIGSACFCYFNC |
| *Cicer arietinum* | XP_027192379.1 | NLCQRKSTTWSGPCLDTGNCKNQCLTVEH--ATFGACHRDGFGFACFCYFNC |
| *Lycium barbarum* | XP_060171152.1 | KLCQRRSKTWSGPCINTGNCSRQCKNQED--ARFGACHRNGFGFACFCYFNC |
| *Trifolium repens* | KAK2352847.1 | KVCQKRSETWTGPCIVTGNCKKQCINVEQ--ATFGACHHQGFGFACFCYFKC |
| *Ipomoea batatas* | GMC58423.1 | KVCQKRSKTWTGPCIITGNCSRQCKNIEG--ATFGACHRSGFGFACFCYFKC |
| *Punica granatum* | XP_031376995.1 | KVCQKRSKTWSGVCLNTGNCNRQCRNWEG--ARSGACHRQGFGFACFCYFKC |
| *Eucalyptus grandis* | XP_039170416.1 | KLCERRSKTWSGFCGNSGNCDRQCKNWEG--ARSGACHAQSLGLACFCYFNC |
| *Olea europaea* | XP_022842068.1 | KVCSRLSRTWSGICLNTGNCDRQCRNWEK--AQHGACHRRGWGFACFCYRQC |
| *Helianthus annus* | KAF5803281.1 | KLCDKRSKTWSGFCEISKNCDKQCRDWEG--AAHGACHRQGLGMACFCYFNC |
| *Solanum verrucosum* | WMV36895.1 | KVCQRRSQTWSGMCINTGNCSRQCKQQED--ARFGACHQNGIGFACFCYFTC |
| *Quillaja saponaria* | KAJ7957773.1 | -ICQRRSKTWSGFCGNSGNCDRQCRNWEG--ATHGACHAQFPGFACFCYFRC |
| *Medicago truncatula* | KEH18159.1 | --CQRKSKTWSGPCLNTANCKNQCISKEP-PATFGACHRDGIGFACFCYFNC |
| *Lathyrus sativus* | CAK8070387.1 | KVCQKRSKTWSGFCANSGNCKRQCIDVES--ATFGACHRQGIGLACFCYFKC |
| *Olea europaea* | CAA2957929.1 | KICSRPSKTWTGPCIKTENCNKQCKNWEE--ARNGACHRSGIGFACFCYFKC |
| *Escallonia herrerae* | KAK3028736.1 | -LCRKRSTTWSGPCLNSNGCKDQCIRLEK-PAVFGACHRQGIGSACFCYYNWC |
| *Abrus precatorius* | XP_027337095.1 | --CQCRSKTWFGPCFDSNGCKNQCINQEQ--ANFGACHRQGIGTACFCYYTC |

**γ-core**

**Supplementary Fig S1.** Amino acid sequence alignment of MtDef5B homologs from different plants. MtDef5B homologs sequences were obtained from NCBI and aligned using Clustal Omega. The *γ*-Core motif is highlighted within the red box and the amino acid differences in the γ-Core motif between the homologs are indicated in blue letters.

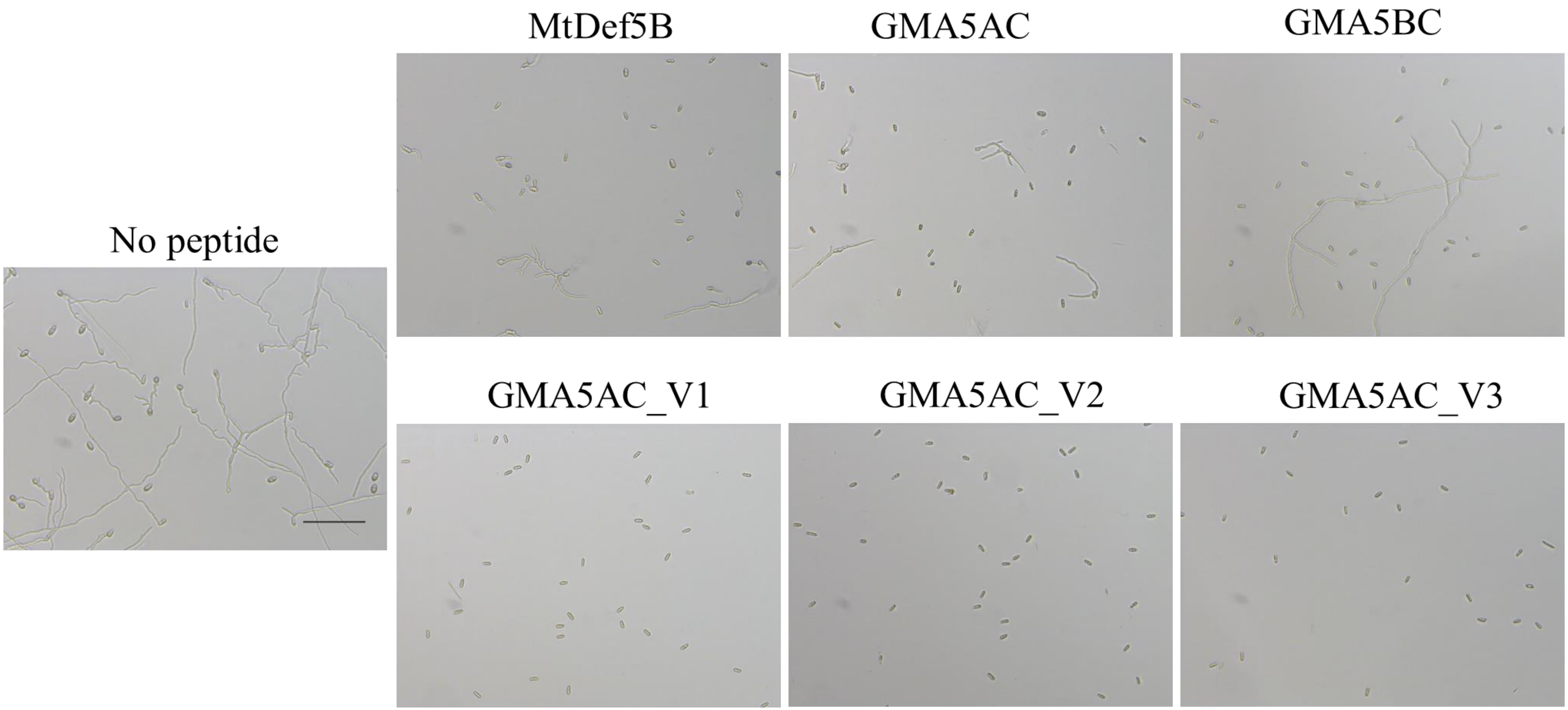


**Supplementary Fig S2.** Representative microscopy images of *Colletotrichum gloeosporioides* treated with 12 μM MtDef5B, GMA5AC, GMA5BC, GMA5C_V1, GMA5C_V2, and GMA5C_V3. Images were taken 24 h after incubation with peptides. Scale bar = 100 μm.

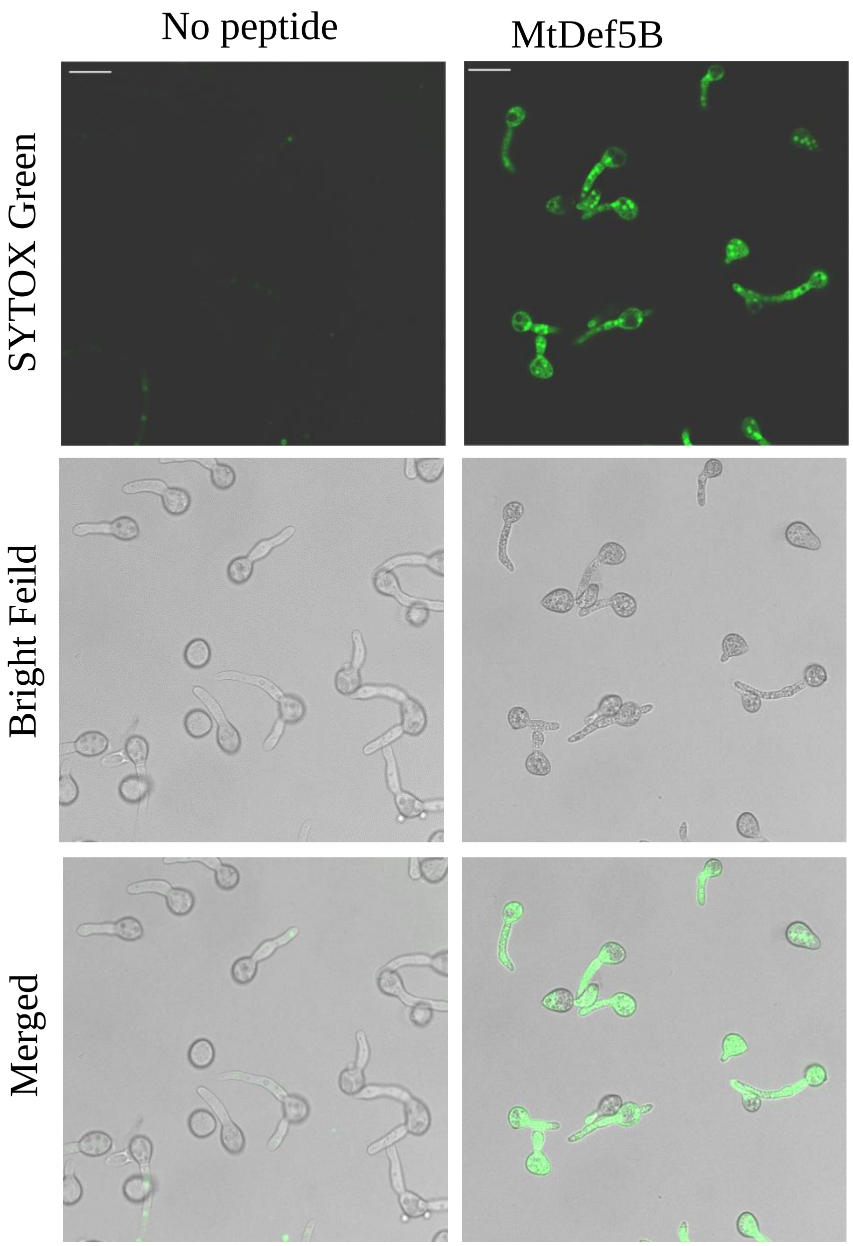


**Supplementary Fig S3.** The MtDef5B permeabilizes the fungal plasma membrane in *Botrytis cinerea* germlings. The fluorescence confocal microscopy images show SYTOX Green (SG) binding to the nuclei following treatment with 0.75 µM of MtDef5B in the presence of 0.5 µM SG after 15 minutes. Scale bar = 20 µm.

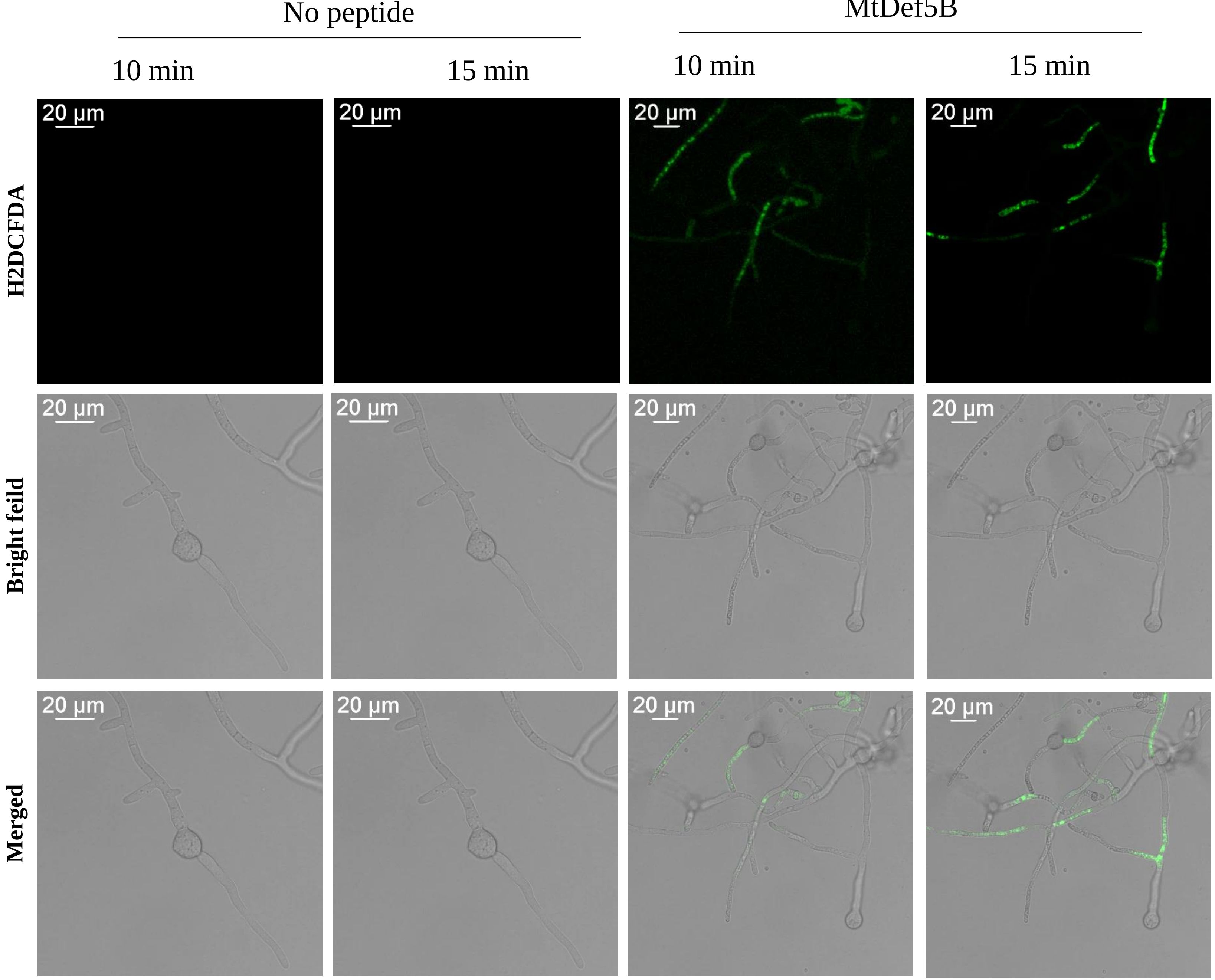


**Supplementary Fig S4**. **MtDef5B induces accumulation of Reactive Oxygen Species (ROS) in *Botrytis cinerea* germlings.** Confocal microscopy images showing ROS production (indicated by green fluorescence from H2DCFDA) in *B. cinerea* germlings after treatment with 0.75 µM MtDef5B for 10 and 15 minutes, Scale bar = 20 µm.

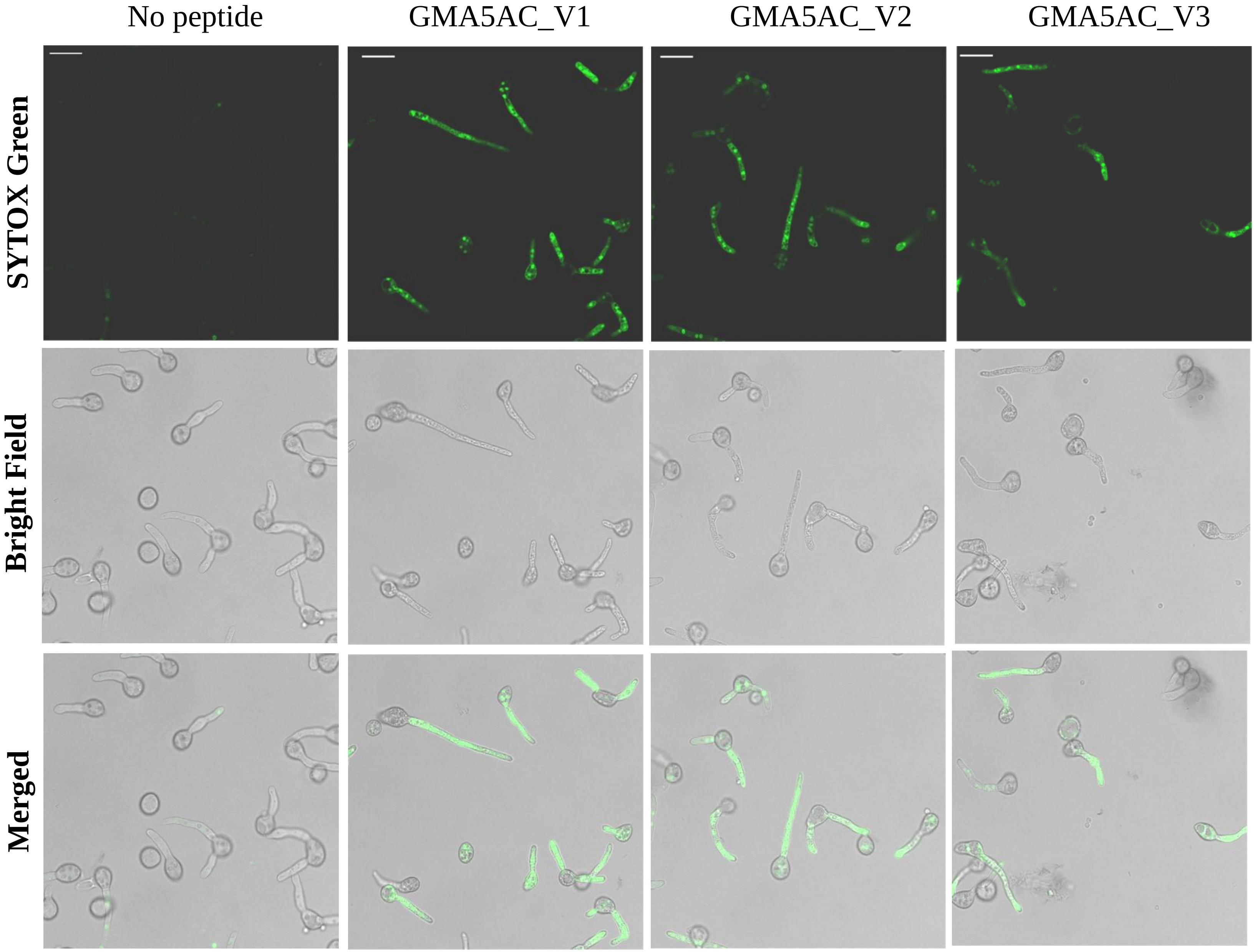


**Supplementary Fig S5.** The short peptide variants of GMA5AC: GMA5AC_V1, GMA5AC_V2, and GMA5AC_V3—permeabilize the fungal plasma membrane in *Botrytis cinerea* germlings. The fluorescence confocal microscopy images show SYTOX Green (SG) binding to the nuclei following treatment with 1.5 μM of the peptides in the presence of 0.5 μM SG after 15 minutes. Scale bar = 20 μm.

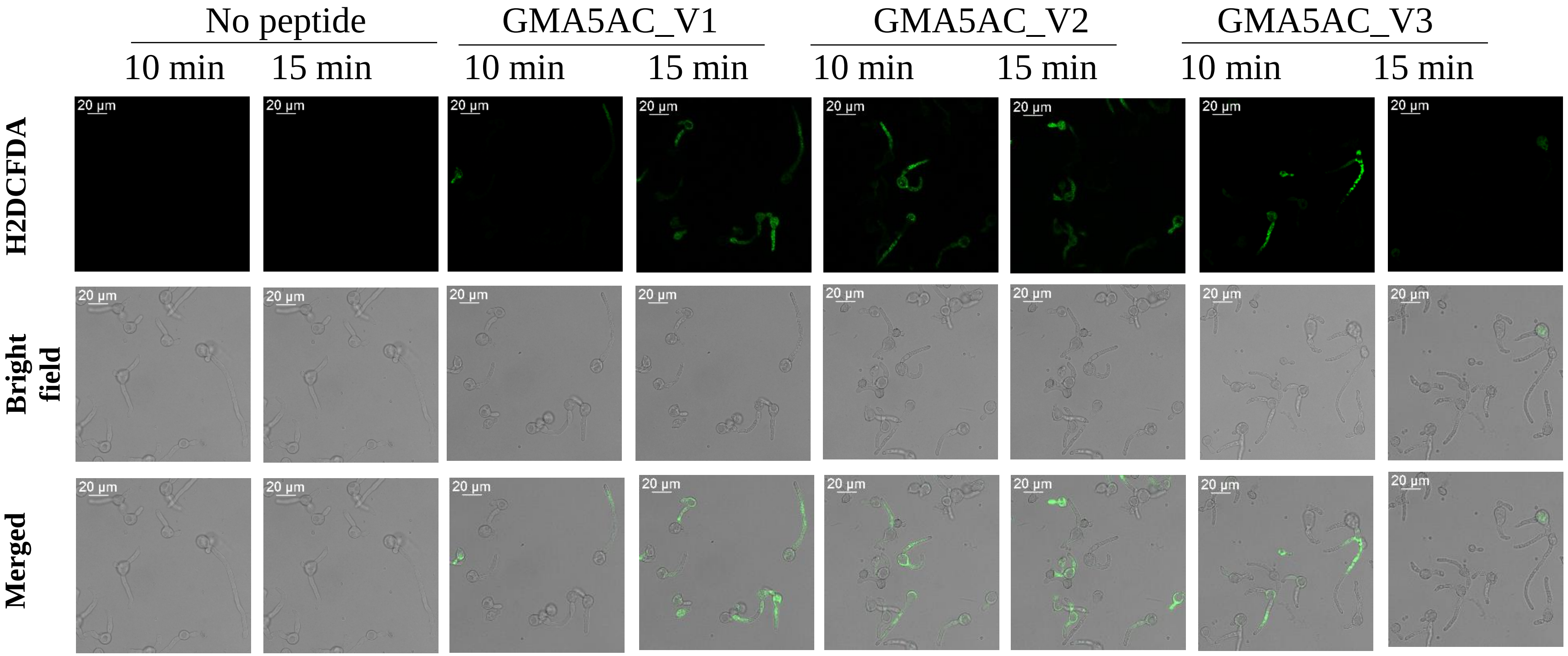


**Supplementary Fig S6. The short peptide variants of GMA5AC: GMA5AC_V1, GMA5AC_V2, and GMA5AC_V3 induces accumulation of Reactive Oxygen Species (ROS) in *Botrytis cinerea* germlings.** The confocal microscopy images showing ROS production (indicated by green fluorescence from H2DCFDA, 10 µM) in *B. cinerea* germlings after treatment with 1.5 µM the peptides for 10 and 15 minutes. Scale bar = 20 µm.

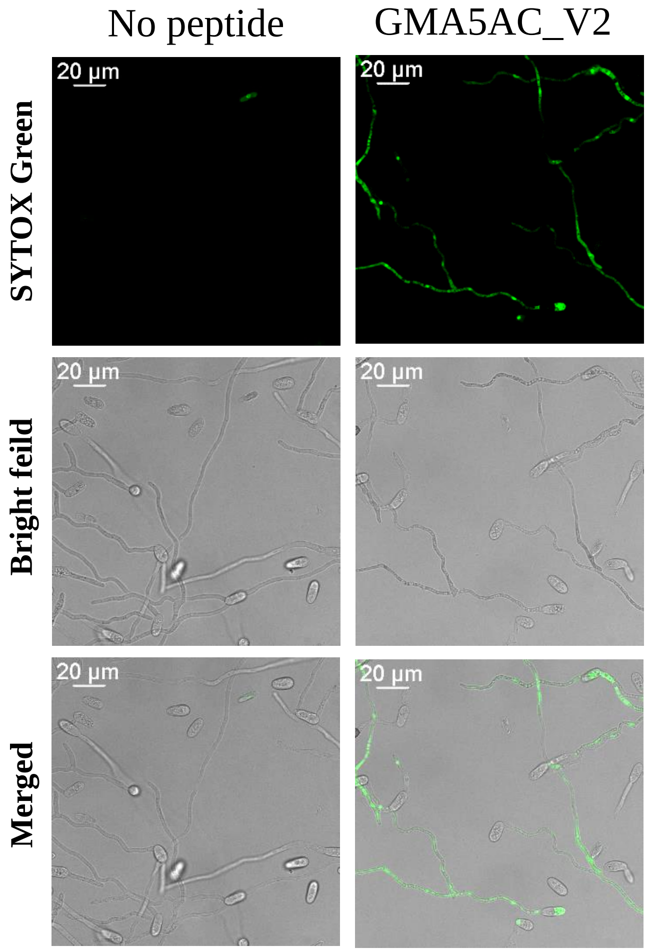


**Supplementary Fig S7.** The GMA5AC_V2 permeabilizes the fungal plasma membrane in *Colletotrichum gloeosporioides* germlings. The fluorescence confocal microscopy images show SYTOX Green (SG) binding to the nuclei following treatment with 3.0 µm of GMA5AC_V2 in the presence of 0.5 µM SG after 15 minutes. Scale bar = 20 µm.

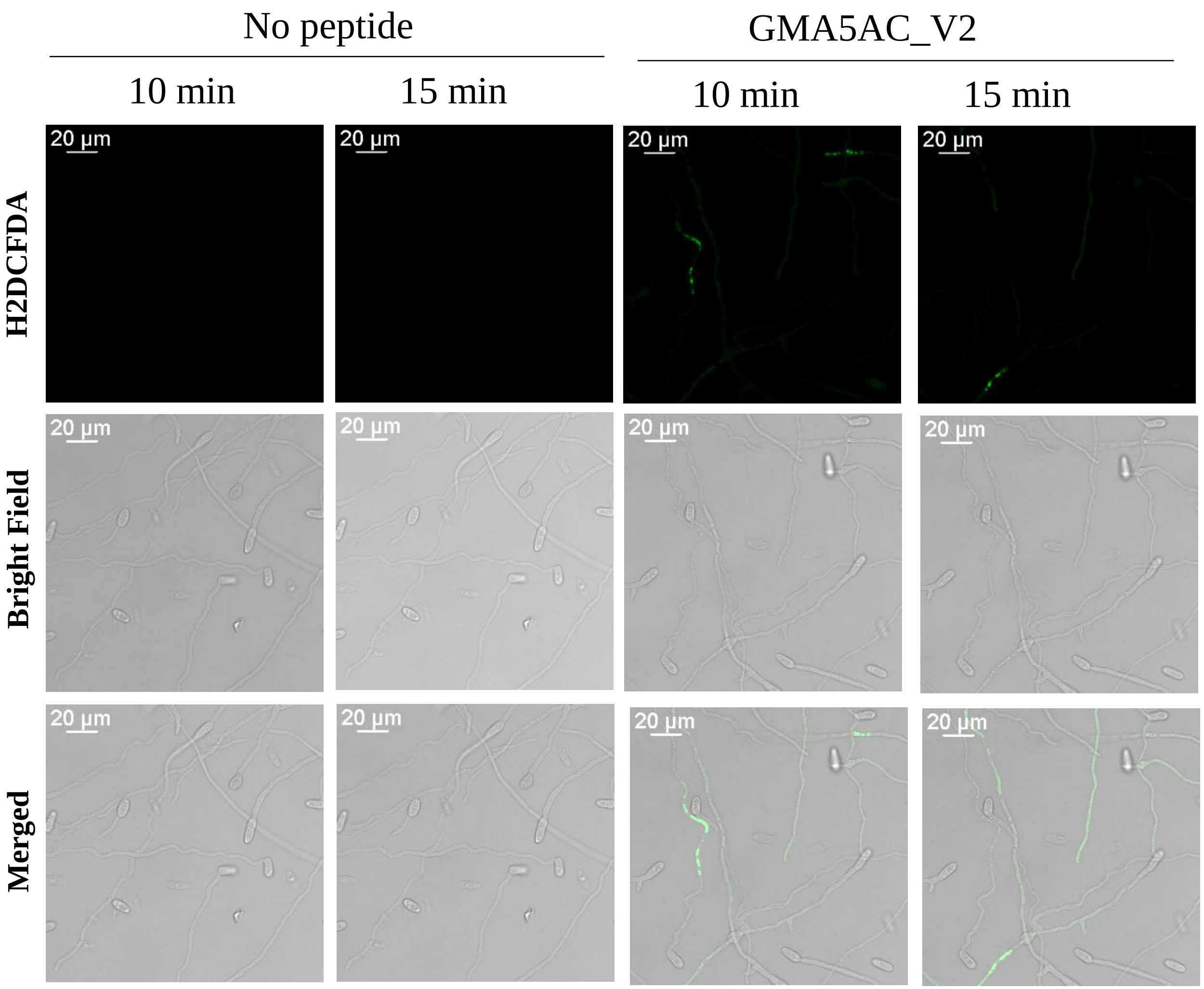


**Supplementary Fig S8. GMA5AC_V2 induces accumulation of Reactive Oxygen Species (ROS) in *Colletotrichum gloeosporioides* germlings.** The confocal microscopy images showing ROS production (indicated by green fluorescence from H2DCFDA, 10 µM) in *C. gloeosporioides* germlings after treatment with 3.0 µM the peptides for 10 and 15 minutes. Scale bar = 20 µm.

**Supplementary Table S1**: Growth media and conditions for culturing the fungi used in this study.

| **Strains** | **Medium for spore production** | **Culture conditions** | **Reference** |
|---|---|---|---|
| *Botrytis cinerea* strain T4 | 20% V8 agar medium | 7-25 days, 25˚C | Lian et al., 2018 |
| *Colletotrichum gloeosporioides* | Potato dextrose agar (PDA) | | MacKenzie et al 2007 |
| *Fusarium virguliforme* | Potato dextrose agar (PDA) | | Veliveli et al 2020 |
| *Fusarium graminerium* | Carboxymethyl cellulose medium (CMC) | | Cappelini & Peterson (1965) |
| *Phytophthora capsica* | Cleared 10% V8-agar (1.5%) medium supplemented with β-sitosterol | | Miller, 1955 |


Cappellini, R. A., and Peterson, J. L. 1965. Macroconidium Formation in Submerged Cultures by a Nonsporulating Strain of *Gibberella Zeae*. Mycologia. 57:962–966

Lian, J., Han, H., Zhao, J., and Li, C. 2018. In-vitro and in-planta *Botrytis cinerea* Inoculation Assays for Tomato. Bio Protoc. 8:e2810

Mackenzie, S. J., Seijo, T. E., Legard, D. E., Timmer, L. W., and Peres, N. A. 2007. Selection for Pathogenicity to Strawberry in Populations of *Colletotrichum gloeosporioides* from Native Plants. Phytopathology. 97:1130–1140

Miller, P. M. 1955. V-8 juice agar as a general purpose medium for fungi and bacteria. Phytopathology 45:461-462

Velivelli, S. L. S., Czymmek, K. J., Li, H., Shaw, J. B., Buchko, G. W., and Shah, D. M. 2020. Antifungal symbiotic peptide NCR044 exhibits unique structure and multifaceted mechanisms of action that confer plant protection. Proc. Natl. Acad. Sci. U. S. A. 117:16043–16054

**Supplementary Table S2**: AlphaFold 2.0 predicted structural confidence and template modeling scores for selected peptides. The table compares various peptides based on the Predicted Local Distance Difference Test (pLDDT) and Predicted Template Modeling (pTM) scores.

| **Peptide** | **Predicted Local Distance Difference Test (pLDDT)** | **Predicted Template Modeling (pTM)** |
|---|---|---|
| MtDef5A | 96 | 0.767 |
| MtDef5B | 95.7 | 0.757 |
| *GMA5AC* | 76.7 | 0.627 |
| GMA5AC_V1 | 58.1 | 0.023 |
| GMA5AC_V2 | 58.0 | 0.229 |
| GMA5AC_V3 | 59.9 | 0.218 |
| GMA5BC | 74.3 | 0.272 |
